# Bypassing the Batch Effects to Improve the Reusability of RAD-seq Data: A Case Study of Recurrent Hybridization in the Japanese *Torreya* Species Complex

**DOI:** 10.64898/2025.12.17.694994

**Authors:** Wenbin Zhou, Shota Sakaguchi

**Affiliations:** Department of Biology, The University of North Carolina at Chapel Hill, Chapel Hill, North Carolina 27599, United States; Graduate School of Human and Environmental Studies, Kyoto University, Yoshida-Nihonmatsu-cho, Sakyo-ku, Kyoto 606-8501, Japan

**Keywords:** genome skimming, gymnosperms, hybridization, phylogenomics, plastome, RAD-seq

## Abstract

Understanding the evolutionary effects of natural hybridization in gymnosperms remains limited due to their large genome sizes and frequent reliance on sparce molecular data. In this study, we reassessed the evolutionary history and taxonomy of the Japanese *Torreya* species complex (*T. nucifera*) by integrating the morphological characteristics, environmental niches, and molecular evidence. We developed RADADOR version 2 to overcome allele dropout (ADO) issues in RAD-seq data sets. This version of RADADOR enabled us to recover ADO loci using low-coverage whole genome sequencing (lc-WGS) data (∼5–10X coverage). We integrated previously generated plastome and RAD-seq data with whole genome data to build robust, genome-wide phylogenies. Subsequently, we employed PhyloNet to detect hybridization events in *Torreya* and utilized a fossilized birth-death model to estimate the formation times of Japanese *Torreya* species. Our phylogenetic results indicated conflicting nuclear and plastome topologies and suggested two historical hybridization events in Japanese *Torreya*. One ancient hybridization between *T. nucifera* and a North American lineage likely contributed to the origin of *T. fruticosa*, while a more recent event may have generated a putative hybrid in regions where their distributions overlap. Integrating the lc-WGS data and RAD-seq data in a reproducible framework revealed the complex reticulate evolution of *Torreya* in Japan and underscores the repeated contribution of inter-specific hybridization to the evolution of gymnosperms.

## Introduction

Hybridization is a fundamental driving force in speciation. It can generate novel genetic diversity and shape biodiversity patterns across different scales (Arnold, 1997; Dowling & DeMarais, 1993; Eckenwalder, 1984; P. R. Grant & Grant, 1992; Rieseberg et al., 2003; Seehausen, 2004). Unlike the conventional view of speciation as a strictly bifurcating process, hybridization often produces complex phylogenetic networks (Soltis & Soltis, 2009). The evolutionary mechanisms that contribute to the occurrence, persistence, and evolution of hybrids remain poorly understood in both plants and animals (Vallejo Marín & Hiscock, 2016). Up to 25% of land plant species and 10% of animal species may involve hybridization events (Genovart, 2009). Approximately 30% to 70% of angiosperms have experienced polyploidy events (Arnold, 1997; Soltis & Soltis, 2009). To date, hybridization and its consequences have been best studied in plants. (Abbott et al., 2013; Arnold, 1997; V. Grant & Grant, 1971; Rieseberg & Carney, 1998). Hybridization events can occur at the same ploidy level (homoploid hybrid speciation), as well as at allopolyploid level (speciation via hybridization and genome doubling) (Soltis & Soltis, 2009; Wu et al., 2022). The best described cases of natural hybridization in plants come from angiosperms and ferns, including *Helianthus* and *Tragopogon* (sunflowers) (Lipman et al., 2013; Rieseberg et al., 2003), *Pteris* (ferns) (Barrington et al., 1989; Soltis & Soltis, 2009), *Quercus* (oak) (Lagache et al., 2013; Petit et al., 2004), and *Mimulus* (Monkey flowers) (Fishman & Willis, 2001). In gymnosperms, hybridization has been reported in wind-pollinated conifers, including *Athrotaxis* (Worth et al., 2016), *Cupressus* (J. Li et al., 2020), *Pinus* (Conkle & Critchfield, 1988; Menon et al., 2021), and *Picea* (Krutovskii & Bergmann, 1995; Tsuda et al., 2016). Hybridizations are not merely transient events, but also have played important roles in gymnosperm speciation and adaptive evolution (Menon et al., 2021; Stull et al., 2021). Given ancient evolutionary history and ecological importance of gymnosperms, understanding the roles of hybridization in gymnosperm evolution and biodiversity may provide critical insights into plant adaptation and speciation mechanisms.

*Torreya* Arn. (Taxaceae) is an example of a woody plant genus with a disjunct distributed in eastern Asia-North America (Donoghue & Smith, 2004; Wen et al., 2010, 2016). The only two species in North America are: *T. taxifolia* Arn., endemic to a limited region of gulf Florida and Georgia in ENA and *T. californica* Torr., endemic to the coastal ranges of California in western North America (Burke, 1975; Hils, 1996; Stalter & Dial, 1984). The current Asian species diversity exceeds that of North American species, including *T. fargesii* Franch, *T. jackii* Chun, *T. grandis* Fortune ex Lindl., *T. yunnanensis* Cheng & Fu (eFloras, 2008), as well as recent discovery on a few potential new Chinese *Torreya* species or varieties, *T. parvifolia* L. and *T. grandis* var. *jiulongshanensis* variety. Zhou et al. (2022) revealed that the disjunct distribution of eastern Asian and North American lineages results from a vicariance event in the Eocene and formation of *T. jackii* is likely caused by the plastid capture events based on RAD-seq data and HybSeq data.

Despite these progress above, an ancient lineage in Japan, *Torreya nucifera* (L.) Siebold et Zucc, that diverged from *T. grandis* in the late Oligocene (Zhou et al., 2022), remains poorly understudied. *Torreya nucifera* (L.) Siebold et Zucc. is a widespread evergreen dioecious conifer species distributed in Japan from northern Honshu to Kyushu and southern areas of the Korean Peninsula and Jeju Island, South Korea (Eckenwalder, 2009; Ohashi 2015; Yamazaki 1995). Ohashi (2015) indicated that it has one variety and four forms, including *T. nucifera* var. *radicans* Nakai in snowy mountains, *T. nucifera* f. *macrosperma* (Miyoshi) Kusaka in Miyagi, Shiga and Mie Prefectures, f. *igaensis* (Doi & Morikawa) Ohwi in Miyagi and Mie Prefectures, f. *nuda* (Miyoshi) Kusaka in Hyogo Prefecture, and f. *sphaerica* (Kimura) Yonekura in Miyagi Prefecture. The taxonomy of variety *T. nucifera* var. *radicans* has been debated for decades. The taxonomic treatment of *T. nucifera* var. *radicans* is commonly accepted based on morphologies (Ōi 1965; Yamazaki 1995); however, combining molecular markers, both Aizawa & Worth (2021), and Ou et al. (2025) considered *T. nucifera* var. *radicans* as a separate species, i.e. *T. fruticosa* Nakai, when using a few plastid DNA (Ou et al., 2025), ITS fragments (Aizawa & Worth, 2021), and a few low copy number genes (Ou et al., 2025). This taxonomic treatment was originally proposed by Nakai (1938) based on their distinct morphologies and habitat, i.e., the unique shrub form and living in snowy mountains. Aizawa & Worth (2021) also emphasized the habitat differentiation between *T. nucifera* and *T. nucifera* var. *radicans*. The latter has narrower distributions in high-evaluation mountains on the side near Sea of Japan with more snow in winters. Although the plastid evidence from Aizawa & Worth (2021) supports the monophyly between *T. nucifera* var. *radicans* and *T. grandis* in China, ITS evidence from Aizawa & Worth (2021) indicated the two varieties of *T. nucifera* may have undergone introgression with *T. nucifera* and *T. grandis*. This introgression was not detected in Ou et al. (2025) due to the absence of closest outgroup *T. grandis* in their study. Most studies only used a few plastid fragments and nuclear genes for phylogenetics (Kou et al., 2017; Mo et al., 2023), likely due to the large genome size in *Torreya* (∼20 Gb in *T. grandis*; Lou et al., 2023), and as such are limited in resolution and power. Without genome-wide molecular evidence, it is unlikely that the long-standing unresolved taxonomic questions about the role of hybridization in *Torreya* can be resolved.

Genome-wide genetic data can provide the genetic signals of hybridization, especially when the events are relative ancient and the signal is subtle and mosaic-like with some regions deriving from one parent, others from the other (Bernhardt et al., 2020; Payseur & Rieseberg, 2016; Taylor & Larson, 2019). Genome-wide markers can minimize the overestimation of one parental lineage’s contribution that can arise from stochastic lineage sorting and the uniparental inheritance of plastid genomes (Currat et al., 2008; Twyford & Ennos, 2012). They also enable the detection of complex reticulate evolutionary patterns shaped by multiple hybridization events and backcrossing. Studies increasingly use using Next-generation sequencing (NGS) technologies to understand the dynamic nature of hybridization and introgression (Hohenlohe et al., 2021; Meleshko et al., 2021; Twyford & Ennos, 2012; Zhou & Xiang, 2022; Zhou et al., 2010). Among all NGS methods, restriction site-associated DNA sequencing (RAD-seq) is one of the most commonplace method for population genomics and phylogenomics studies (Hohenlohe et al., 2021; Twyford & Ennos, 2012). It can generate many thousands of loci across different populations and closely related taxa without any prior genomic reference (Andrews et al., 2016). While more expensive, many studies have advocated that low-coverage whole genome sequencing data (lc-WGS, ∼5x to 10x coverages) can get accurate genotyping data as an alternative to RAD-seq (Buerkle & Gompert, 2013; De La Harpe et al., 2017; Kumar et al., 2021).

However, both methods encounter some disadvantages for phylogenetics and population genetics. Regarding whole genome sequencing, it is still too costly for population genetics projects with many accessions involved, especially for non-model species with large genome sizes. Additionally, it requires a well assembled genome to obtained informative SNPs, which remains a challenge for many non-model species. In terms of RAD-seq, the primary disadvantage is allele dropout (ADO), which is usually caused by either mutations and evolutionary changes in restriction sites among taxa or inconsistency of library construction (e.g. diverse DNA qualities, different restriction enzymes, PCR cycles, and size selection of libraries). This can ultimately lead to unpredicted missing data in down-stream analyses for both phylogenetics and population genetics (Andrews et al., 2016; Davey et al., 2013; Gautier et al., 2013; Leaché & Oaks, 2017; Mastretta-Yanes et al., 2015). Unlike traditional barcoding method (*rbcL* and *trnL-trnF*) and modern target enrichment methods (Johnson et al., 2019; Mamanova et al., 2010; Zhou & Xiang, 2022), RAD-seq is not comparable among different sets of experiments because the ADO is particularly profound (Andrews et al., 2016). To mitigate ADO, rigorous standardization is essential. Namely, all samples should be processed simultaneously and using a same size selection strategy to reduce the batch error and maximize the number of shared alleles for downstream analyses. Even minor variance in wet-lab processes can lead to non-overlapping loci among different datasets (i.e. batch effects), which would largely reduce the number of shared RAD-seq loci if the experiments were conducted at different times and combined later (Malinsky et al., 2018). For example, if a critical population was newly discovered, generating a new RAD-seq dataset might not be compatible with the published RAD-seq dataset due to ADO-driven data incompatibility. These limitations in both WGS and RAD-seq can impede the long-term research continuity. To address this issue and make the ongoing RAD-seq projects compatible, we propose using low-coverage whole genome sequencing (lc-WGS) data (∼5x to ∼10x) as an efficient alternative to overcome ADO challenges for newly collected populations and make previous RAD-seq data reusable—even when a reference genome is unavailable. By leveraging raw WGS reads, our newly developed RADODOR v.2 which adopts the main methods from RADODOR (Zhou et al., 2022) and GeneMiner (Xie et al., 2024) to identify and extract loci shared across populations, enabling consistent variant calling without the batch effects inherent to RAD-seq.

Here we successfully resolved the phylogeny of two Japanese *Torreya* taxa by integrating environmental factors (i.e., temperature, precipitation, altitude, and snow coverage), genome-wide molecular data, and plastome evidence to better understand how potential ancestral hybridization events shaped this clade. Specifically, we: (1) compare niche differentiation and structural differentiation of plastomes between two *Torreya* taxa in Japan; (2) develop a RADADOR version 2—a unique tool for extract all corresponding published RAD-seq loci from newly whole genome sequencing data, (3) reconstruct a robust phylogeny of *Torreya* species based on both genome wide nuclear data and plastomic data to clarify the taxonomic ranks of Japanese *Torreya* complex; and (4) reveal hybridization events to elucidate the evolutionary history of Japanese *Torreya*. Based on the evidence from morphologu, ecological niche, and genome-wide molecular data in this study, we hereafter refer to *T. nucifera* var. *radicans* as *T. fruticosa* to reflect their revised taxonomic status and avoid confusion.

## 2. Methods and Materials

### 2.1 Sample Collection and DNA Extraction

We collected 16 *T. fruticosa* samples from two locations including Furuya, Kutsuki-mura Shiga Prefecture and Inami, Mie Prefecture, Japan (Table 1). DNA was extracted by a modified CTAB method following Doyle (1991) and Zhou *et al*. (2022). Due to the large genome size of *Torreya* (∼20 Gb in *Torreya grandis* by Lou *et al*., 2023), we first amplify three gene regions, including two plastid and one ITS fragments (Aizawa & Worth, 2021), to verify the plastid haplotypes and the ribosome types. Specifically, we designed *rbcL* (∼550 bp) forward primer: GCATTCCGAGTCACTCCTCA and reverse primer: AAACGATCTCTCCAGCGCAT; *trnL-trnF* (∼440 bp) forward primer: TCCCCAGGTTTATGTCCGAA and reverse primer: AACTGGTGGCACGAGGATTT; and *ITS* (∼800 bp) forward primer: TGTTGGTTTTGAGCCCACGA and reverse primer: TGCTTAAACTCAGCGGGTGT, based on Primer3 (Untergasser et al., 2012). Then we selected six out of 16 *T. fruiticosa* individuals for low-coverage whole genome sequencing (∼100 Gb per individual) based on their distinct genotypes, including both C1 and C2 haplotypes (Table 1), to avoid sequencing the samples with similar genetic backgrounds. Library were constructed using MGIEasy PCR-Free DNA Library Prep set (MGI Tech). Then the four samples were sent for DNBSEQ-T7 sequencer at UNC-Chapel Hill and two samples were sent to DNBSEQ-G400 sequencer in Japan.

### 2.2 Niche Differentiation

To investigate niche differentiation between *T. nucifera* and *T. fruticosa*, we first downloaded all occurrence records for both species from GBIF (*T. nucifera*: GBIF.org [25 February 2025] https://doi.org/10.15468/dl.eeram8; *T. fruticosa*: GBIF.org [25 February 2025] https://doi.org/10.15468/dl.zc5czd). After manual validation, we only kept occurrences from Japan and excluded *T. fruticosa* records from the *T. nucifera* dataset and any misidentified or human cultivated records. For each validated occurrence, we extracted environmental data from 19 bioclimatic variables (including temperature, precipitation), one altitude available in WorldClim (https://www.worldclim.org/data/bioclim.html), and a custom layer of monthly maximum snow depth across Japan from (https://nlftp.mlit.go.jp/ksj/gml/datalist/KsjTmplt-G02-v3_0.html) using the ‘raster’ package in R (version 4.4.0) (Hijmans et al., 2015). Then, we applied Student’s t-tests to quantify the niche differentiation between the two taxa. All results were visualized in boxplots by ggplot2 in R (version 4.4.0) (Wickham, 2011).

### 2.3 Plastome Assembly and Annotation

We conducted getOrganelle (Jin et al., 2020) to assemble the plastome for all six *T. fruticosa* individuals using ∼10 Gb paired-end reads. Assembled plastomes of all six individuals were visualized by Bandage (Wick et al., 2015). Then, all plastomes were annotated using GeSeq online (Tillich et al., 2017). Then we manually checked the start and stop codons of each annotated gene using Geneious version 2024 (Kearse et al., 2012) according to the annotation results from published *T. nucifera* (MK978775; Shin *et al*., 2019). Finally, we draw circular representations using CIRCOLETTO (Darzentas, 2010). To detect rearrangement events within *Torreya*, we used progressive Mauve (Darling et al., 2004) to examine the arrangements of locally colinear blocks (LCBs) based on the alignments of *T. grandis*, *T. fargesii*, *T. yunnanensis*, *T. nucifera*, and our newly assembled *T. fruiticosa*.

### 2.4 Genome-Wide Nuclear Data

To obtain shared RAD-seq loci from lc-WGS data for newly collected *T. fruticosa* individuals for phylogenetic analyses, we developed RADADOR version 2. Briefly, we incorporated GeneMiner (Xie et al., 2024), a k-mer based method for target gene assembly, to filter all reads from lc-WGS data based on previous sequenced RAD-seq loci (Figure 1). This version of pipeline can take output files from either ipyrad (Eaton & Overcast, 2020) or RADADOR (Zhou et al., 2022). Subsequently, it conducts GeneMiner (Xie et al., 2024) using lc-WGS clean data to filter all reads and assemble all loci matching to RAD-seq loci. To make sure all assembly results are reliable, RADADOR version 2 can filter and keep all loci covered by more than four reads. Finally, it merge all corresponding loci from each newly sequenced individual to each RAD-seq locus, followed by MAFFT (Katoh & Standley, 2013) and TrimAL (Capella-Gutiérrez et al., 2009). Pipeline is open access and available on GitHub (https://github.com/Bean061/RAD_Allele_Dropout_Remedy2).

**Figure 1.**
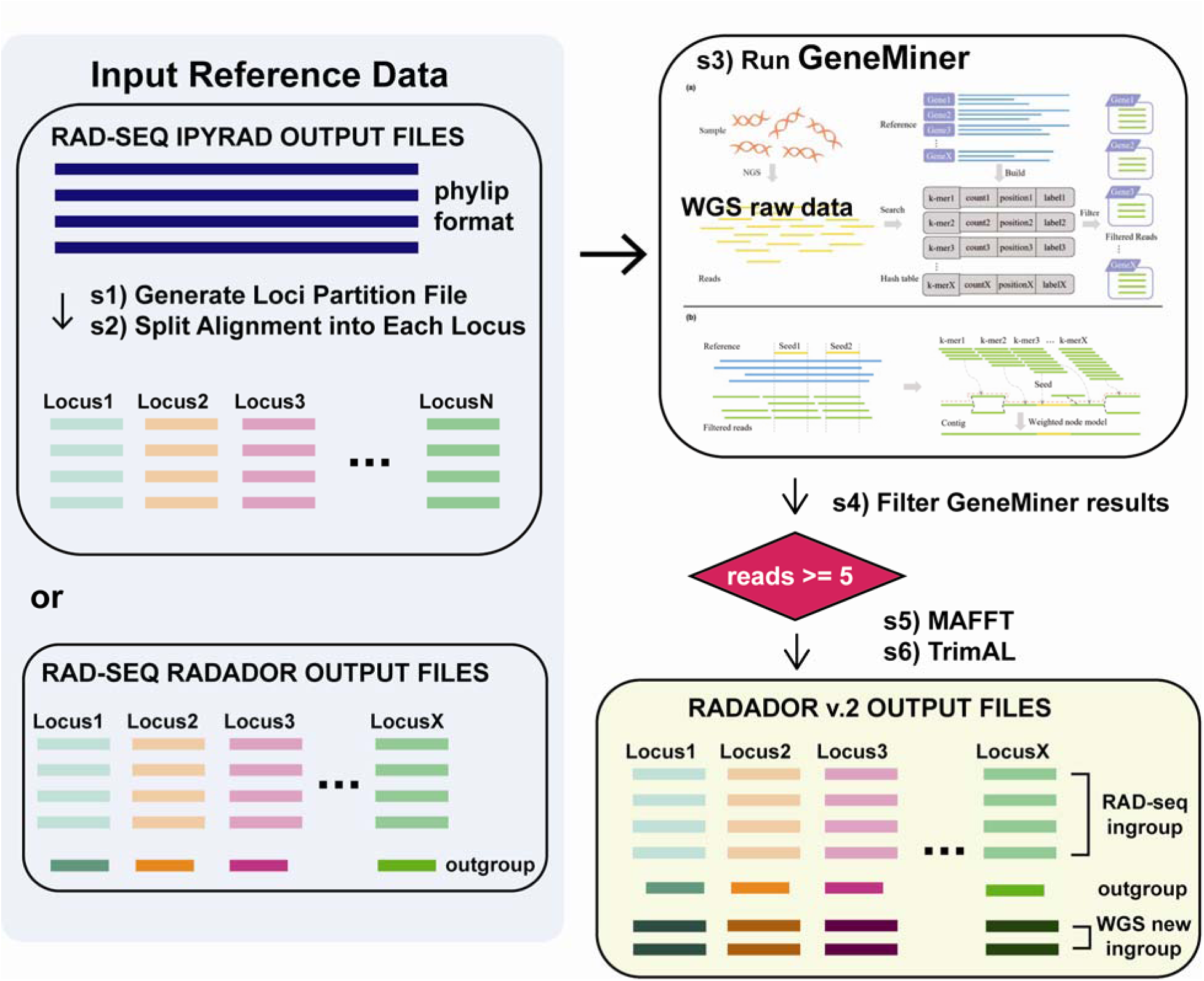
RADADOR version 2 pipeline for Locus Recovery from Whole Genome Sequencing Data. Input: output loci from ipyrad or RADADOR version 1 and trimmed whole genome sequencing data. Steps: it runs GeneMiner to identify and assemble WGS reads matching RAD-seq loci using k-mer alignment. Output: it generates the final loci files with newly sequenced individuals.

### 2.5 Phylogeny

We carried out phylogenetic analyses using two datasets, i.e., genome-wide nuclear loci from RAD-seq and lc-WGS, and plastomic data from lc-WGS, to evaluate the evolutionary consistency between nuclear and plastid data. To obtain the *Torreya* nuclear phylogeny with *T. fruticosa* and its putative hybrid, we first obtained RAD-seq M50 loci of *Torreya* from Zhou et al. (2022) and used the lc-WGS clean data of six individuals of *T. fruticosa* trimmed by fastp (Chen et al., 2018) using default parameters to retrieved the shared RAD-seq data via RADADOR v. 2. After we obtain all newly combined RAD-seq loci alignments, we applied catfasta2phyml.pl script (https://github.com/nylander/catfasta2phyml) to concatenate all loci and applied IQ-TREE 2 (Minh et al., 2020) to generate a phylogeny with “TESTNEW” model and partitioned by each locus. To reconstruct possible network-like evolutionary relationships within the *T. nucifera* species complex, we carried out neighbor net analysis in SplitTree4 (Huson & Bryant, 2006) with combined RAD-seq loci from RADADOR v. 2.

To build the plastid phylogeny, we utilized the plastome matrix from Hyb-Seq data (Zhou et al., 2022) as backbone, then add the newly assembled six plastomes *T. fruticusa* using MAFFT “-add” function. Then, we conducted IQ-TREE 2 (Minh et al., 2020) to reconstruct the plastid phylogeny with “TESTNEW” model without any partition. Both genome-wide nuclear and plastomic matrices were deposited in Dryad (DOI: 10.5061/dryad.59zw3r2n1).

### 2.6 Hybridization Event and Divergence Time Estimation

To disclose the hybridization history of *T. nucifera* complex, we employed PhyloNet (Than et al., 2008) -- a gene tree-based coalescent method -- to detect potential hybridization events. Given that the ancient divergence time of *Torreya* (∼180 Mya; Zhou et al., 2022) may result in substitution saturation at neutral sites and various substitution rate among lineages, the ABBA-BABA test analyses could be skewed (Frankel & Ané, 2023) and, therefore, was abandoned here. Specifically, we implemented the Maximum Pseudo-likelihood (MPL) approach in PhyloNet to test hypotheses with one versus two hybridization events in *Torreya*. Our analyses consisted of two phases. First, we treated all six *T. fruticosa* (including the potential hybrid) as a monophyletic group. Subsequently, we divided *T. fruticosa* into two distinct lineages, including one lineage representing the potential hybrid and the other lineage consisting of all remaining *T. fruticosa* specimens.

To estimate the divergence time of *T. nucifera* complex, we incorporated fossil morphologies and applied fossilized birth and death (FBD) model following Zhou et al. (2022) using BEAST v2.6.7 (Bouckaert et al., 2014) to estimate the divergence time. We added the morphology including the leaf lengths, seed sizes, and habits for *T. fruticosa* (supplementary Table S2) to the trait matrix from Zhou et al. (2022). The FBD data matrix was deposited in Dryad (DOI: 10.5061/dryad.59zw3r2n1).

## 3. Results and Discussion

### 3.1 Niche differentiation

Japanese *Torreya* species are differentiated in niche. The student t-tests on two *Torreya* taxa (Fig. 2A) indicated a significant (*P* values < 0.05) niche differentiation between *T. fruticosa* and *T. nucifera* in terms of multiple factors, including 20 out of 21 environmental factors (Fig. 2B and Fig. S1). Overall, we detected lower temperature (e.g. Annual Mean Temperature, Min Temperature of Coldest Month, Mean Temperature of Coldest Quater), more precipitation (e.g. Precipitation in Driest Month, Precipitation in Driest Quarter, Precipitation of Coldest Quarter), and thicker snow depths in *T. fruticosa* compared to *T. nucifera* (Fig. S1).

**Figure 2.**
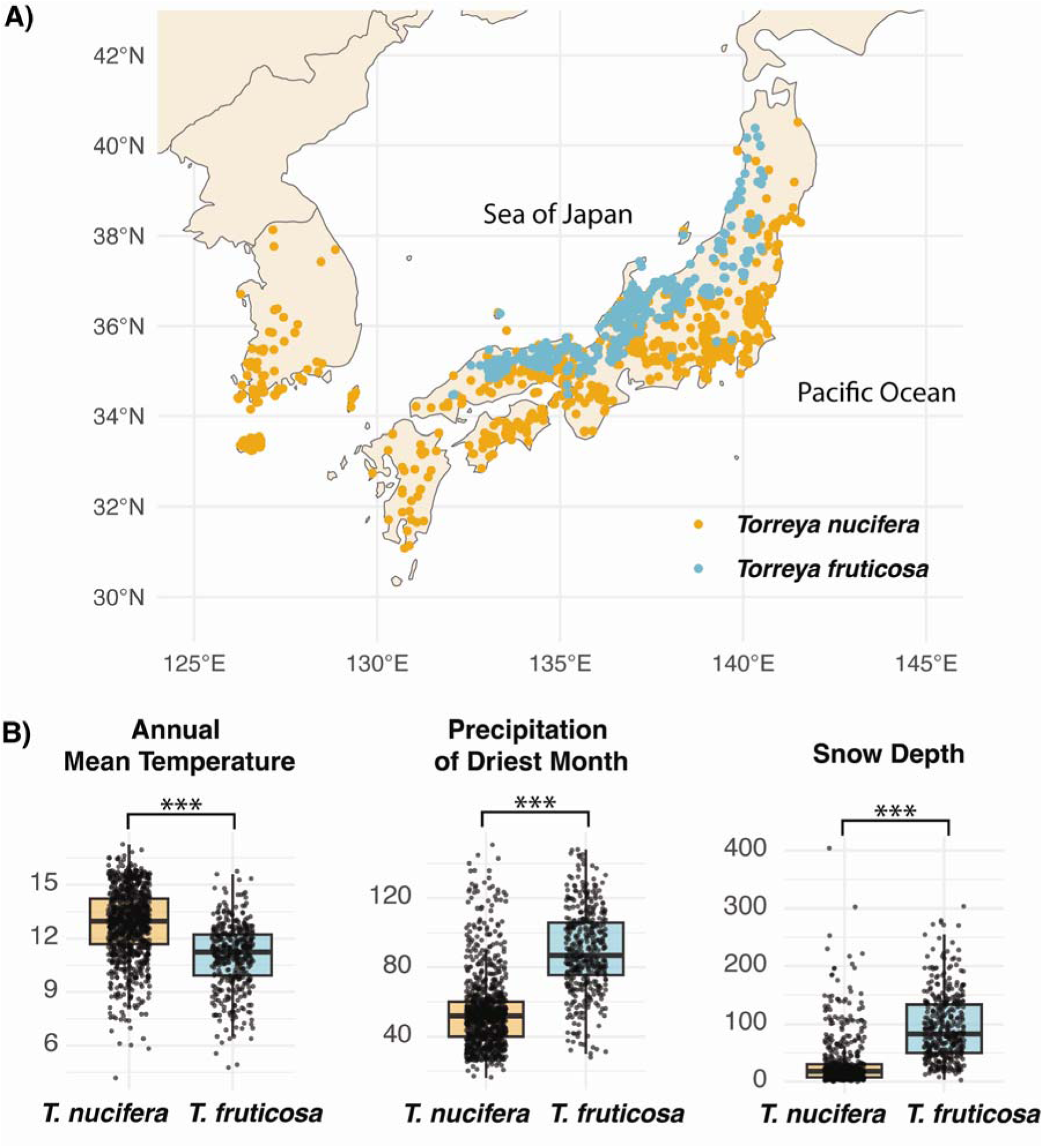
The niches and distributions of two Japanese *Torreya* taxa are distinct. Overall, *T. fruticosa* distributes in the high-elevation mountains on the side near Sea of Japan with lower annual temperature, higher precipitation of driest month and more intense snow in winters, while *T. nucifera* has a wider distribution towards the side near Pacific Ocean. A) Distribution map of two taxa. B) Three major differentiated environmental factors of two taxa. See all comparisons of 21 environmental factors in Fig. S1.

### 3.2 Plastome assembly and annotation

We successfully assembled six plastomes of *T. fruticosa* from the lc-WGS data. All *T. fruiticosa* individuals, except putative hybrid B, had plastomes with an average length of 136,670 bp (Table S1). All six individuals of *T. fruticosa* exhibited an atypical looped plastome structure with several repeats that generated alternative paths in the assembly graph (Fig. 3A), which could not be resolved by using short reads only. In total, we annotated 121 genes for all *T. fruticosa* individuals and its putative hybrid (Fig. 3B). The annotation of these six *Torreya* plastomes revealed 82 unique protein coding genes (PCGs), 30 tRNA genes, and four rRNA genes. Compared to the plastome of *T. nucifera*, both *T. fruticosa* and its putative hybrid retain an identical *trnQ*-inverted repeat (278 bp) but lack the typical long inverted repeat (IR) region in angiosperms (Fig. 3B). The putative hybrid has highly similar GC content to *T. nucifera* and *T. fruticosa* (0.1% less, Fig. 3B). Although *T. nucifera* and *T. fruticosa* shared extremely conservative in gene contents and plastome structure, a few genes from *T. fruticosa* have slight mutations, including an 18-bp deletion in *rpoC2*, 6-bp insertion in *infA*, 12-bp insertion in *rps3*, 36-bp insertion in *ycf2*. In *T. nucifera*, we found *rps2* has 30-bp insertion compared to *T. fruticosa*. The *ycf1* is the most diverged gene which involves at least six insertions/deletions, leading to 2350 aa in *T. nucifera*, 2372 aa in *T. fruticosa*, and 2363 aa in the putative hybrid B (Fig. S4). Besides *ycf1* gene, we detected *accD* gene includes multiple substantial mutations. Specifically, three major insertions/mutation among *T. nucifera*, *T. fruticosa*, and its putative hybrid were found, leading to 879 aa, 823 aa, and 887 aa, respectively (Fig. 3C).

**Figure 3.**
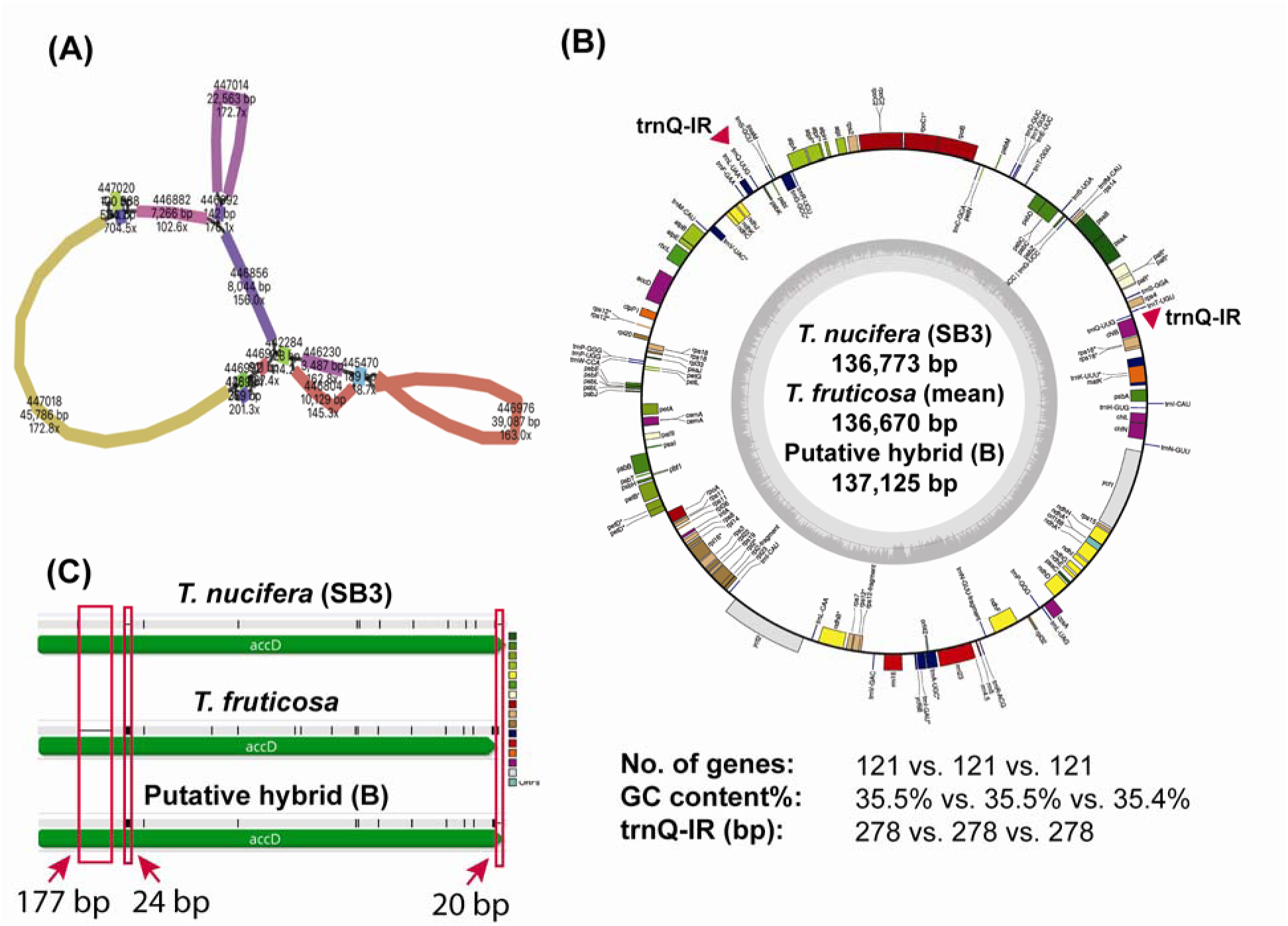
Plastome assembly and annotation of *Torreya nucifera* species complex reveals conservation of plastomic architectures. A) Bandage visualization result indicates the plastome of *T. fruticosa* consists of many long repeats, leading to unresolved assembly paths. B) all plastomes of *T. nucifera* species complex shared a total of 121 genes and similar GC contents. No typical long-IR regions but 278-bp long trnQ-IR regions were found in all three taxa (indicated by the red triangles). (C) The *accD* gene mutation was found in *T. fruticosa* caused by a 177-bp deletion, two insertions (24 bp insertion and 20 bp insertion), which were indicated by the red arrows. Putative hybrid (B) has an admixture structure of *accD* gene with *T. nucifera* (177-bp insertion, and 20-bp deletion) and *T. fruticosa* (24-bp insertion).

### 3.3 Phylogeny conflict between nuclear genome and plastome

We successfully obtained an average of 1801 out of 2731 loci in six newly sequenced *T. fruticosa* using ∼5x lc-WGS data (Table S1) for the downstream phylogenetic analysis. The nuclear phylogeny revealed that *T. fruticosa* is a monophyletic group sister to *T. nucifera* with the putative hybrid B at the base of *T. fruticosa* clade (Fig. 4A). The plastid phylogeny results indicated all *T. fruticosa* except the putative Hybrid B forms a monophyletic group sister to *T. grandis* (Fig. 4B) and the putative Hybrid B has nearly identical plastome genome with *T. nucifera* (Fig. 4B). Two *T. fruticosa* individuals from Aizawa & Worth (2021) belonged to *T. fruticosa* clade (Fig. 4B). SplitTree result from genome-wide nuclear data also supported a monophyly of five *T. fruticosa* individuals and revealed Hybrid B nested in between clades of *T. fruticosa* and *T. nucifera* (Fig. 4C).

**Figure 4.**
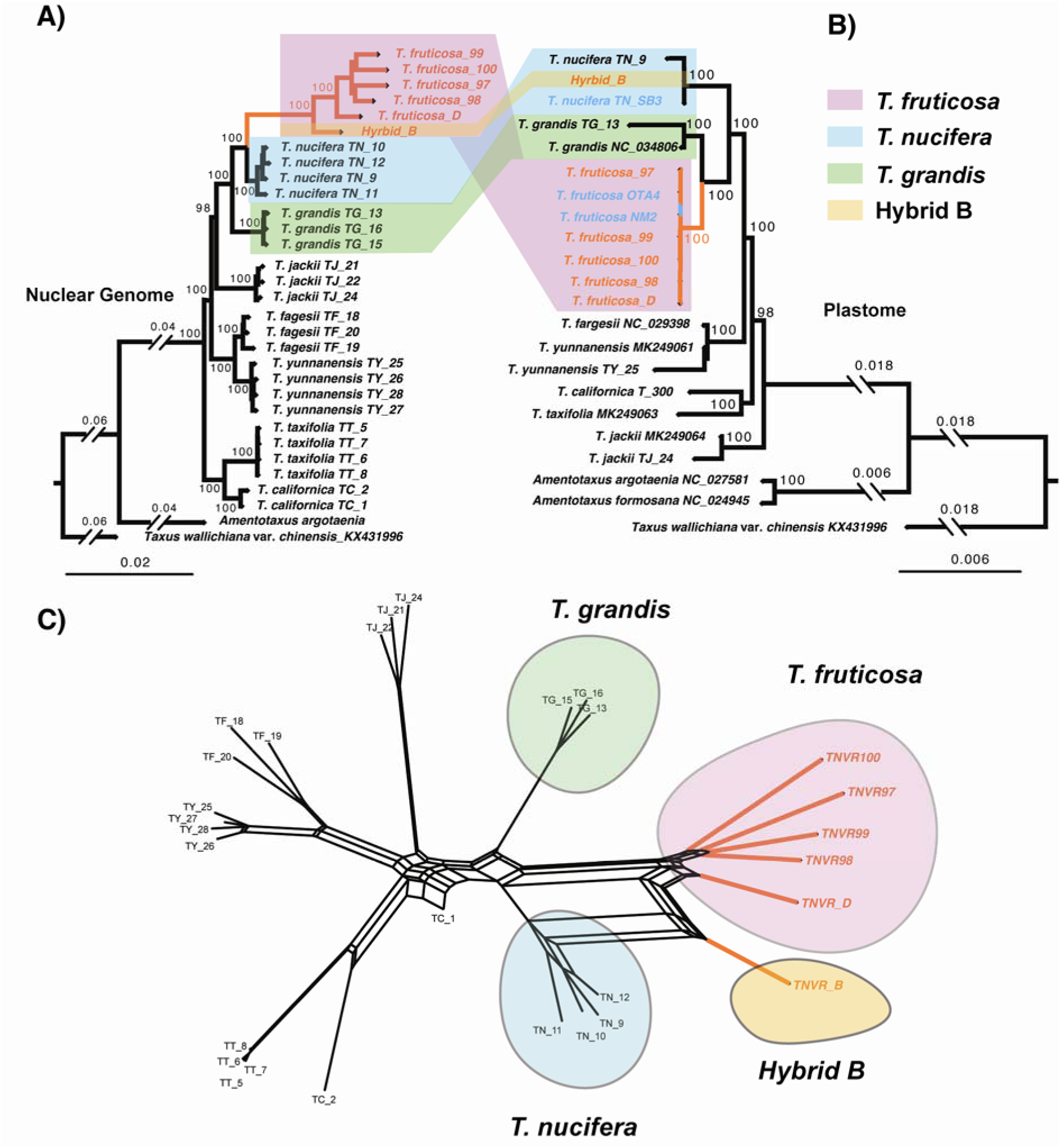
Phylogenetic analyses suggest *Torreya fruticosa* forms a monophylic group and highlights the conflicting placement of *T. fruticosa* and the putative Hybrid B, indicating potential ancient hybridization events. A) phylogenetic result based on combined RAD-seq loci and low-coverage whole genome sequencing data indicates *T. fruticosa* is sister to *T. nucifera* and putative Hybrid B has similar genetic background with *T. fruticosa*. B) Plastid phylogenetic result supports *T. fruticosa* is sister to *T. grandis* and Hybrid B is close to *T. nucifera*. Support values at nodes are UFBS values from IQ-TREE. C) SpilitTree result suggests that *T. fruticosa* forms its own clade and the potential hybridization event led to Hybrid individual B. The orange tips were newly sequenced in our study, black tips were from Zhou et al. (2022), while blue tips were from Aizawa & Worth (2021). The letter “T.” is abbreviation for *Torreya*.

### 3.4 Hybridization Events and Divergence Time

The phyloNetwork analysis indicated two potential hybridization events in *Torreya* (Fig. 5A). One ancient hybridization event (H1) occurred between an American species (likely *T. californica*) and *T. nucifera* to form the *T. fruticosa*. A total of 91% genome was shared between *T. nucifera* and *T. fruticosa* (Fig. 5A). This event was also detected when assuming only one hypothetic hybridization event in the history (Fig. S3). A more recent hybridization event (H2) appeared between the *T. nucifera* and *T. fruticosa*, forming a potential Hybrid B with 86% genomes shared with *T. fruticosa* (Fig. 5A). Our Fossilized Birth-Death result suggested that *T. fruticosa* was formed around 21.7 Mya (Fig. 5B).

**Figure 5.**
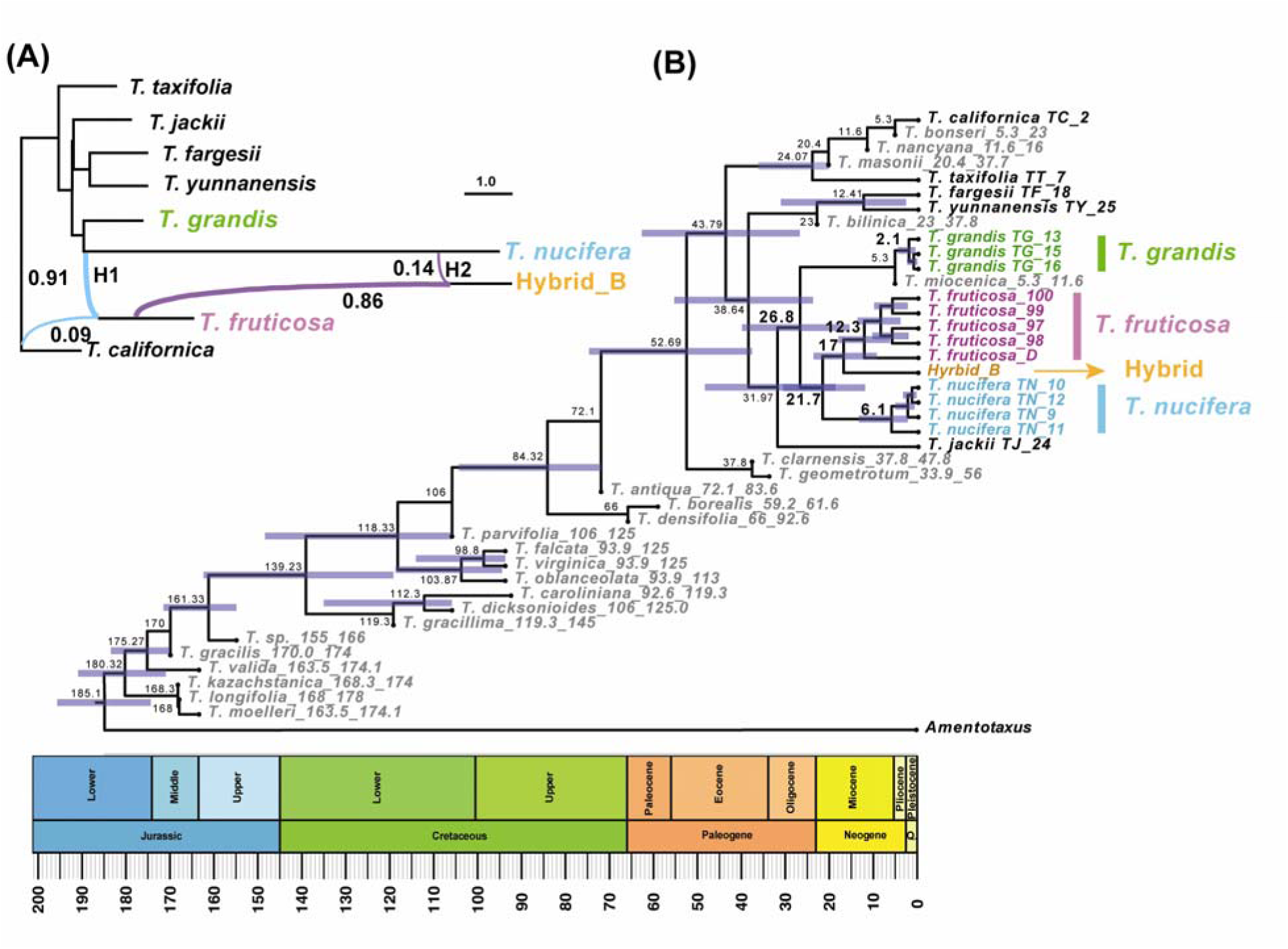
Divergence time and network analyses support two ancient hybridization events in *Torreya*. (A) phyloNetwork result indicates two hybridization events formed *T. fruiticosa* and the putative hybrid. (B) Fossilized Birth and Death model reveals the divergence time between *T. fruticosa* and *T. nucifera* may trace back to 21.7 Mya. The gray tips indicate the extinct species of *Torreya* in panel B.

## 4. Discussion

### 4.1 Taxonomic Treatment and Formation of *T. fruticosa* and Potential Hybrid

*T. fruticosa* prefers a colder, wetter, and higher altitude habitat compared to *T. nucifera* (Fig. 2). Notably, the difference in snow depth supports the idea that the shrubby and compact growth form of *T. fruticosa* was selected in response to the heavy snowfall in regions facing the Sea of Japan, where snowfall can be exceptionally extreme (Fig. 2). Despite the close evolutionary relationship between *T. nucifera* and *T. fruticosa* (Fig. 4, Fig. S2), the distinct morphology (Tree vs. Shrub), habitats (Fig. 2, Fig. S1), and ancient divergence time of *T. fruticosa* (Fig. 5) support its recognition as a separate species. Given all evidence above, we propose that formally adopting the name *Torreya fruticosa* Nakai for *T. nucifera* var. *radicans*. Our results agreed with the taxonomic conclusion from Aizawa & Worth (2021) and Ou et al. (2025), and further fully resolve the phylogenetic placement of *T. fruticosa* and relationship with *T. nucifera* and *T. grandis* via genome-wide genetic data. Our finding also helps explain the *T. fruticosa* accessions (To35 and To36) from Atalanta Botanical Garden might be misidentified as *T. nucifera* (Mo et al., 2023), which shows identical nrDNA cistron with *T. nucifera* but closer plastid relationship to *T. grandis*.

Besides the taxonomic ranking of *T. fruticosa*, our results indicate that a putative hybrid zone in Inami, Mie Prefecture, where harbors both *T. nucifera* and *T. fruticosa* species. The morphological traits of some collections from Inami, Mie Prefecture, are with intermediate tree heights and intermediate leave lengths (Table S2). Although not all 12 individuals from Inami, Mie Prefecture were sequenced by lc-WGS, we found five out of them possessed C1 cpDNA haplotype by using two plastid regions (*trnL-trnF* and partial *rbcL* gene region), which are identical to *T. nucifera* but distinguished from typical *T. fruticosa* clade (Table S1). Assuming the likely maternal chloroplast inheritance in *Torreya* (Buchholz, 1940; Mo et al., 2023), we propose that the formation of *T. fruticosa* and the putative hybrid resulted from two hybridization events (Fig. 6): including one ancient hybridization event via predominant gene introgression from *T. nucifera* occurred in the late Oligocene (∼21.7 Mya in Fig. 5), while another hybridization event via major gene introgression from *T. fruticosa* occurred more recent (Fig. 5). The putative hybrid *Torreya* was recently generated through introgression between the two species. Since these plants may represent a transient entity, we do not consider them as a distinct taxon yet. The final validation of hybrid zone should be carefully accessed by further sampling across additional hybrid zone and with more genome wide molecular data, including estimating their formation time and population structures, which could also resolve the taxonomic placement of three other forms of *T. nucifera* from the Wild Flowers of Japan (Ohashi, 2015), including *T. nucifera* f. *macrosperma* (Miyoshi) Kusaka in Miyagi, Shiga and Mie Prefectures, f. *igaensis* (Doi & Morikawa) Ohwi in Miyagi and Mie Prefectures, f. *nuda* (Miyoshi) Kusaka in Hyogo Prefecture, and f. *sphaerica* (Kimura) Yonekura in Miyagi Prefecture.

**Figure 6.**
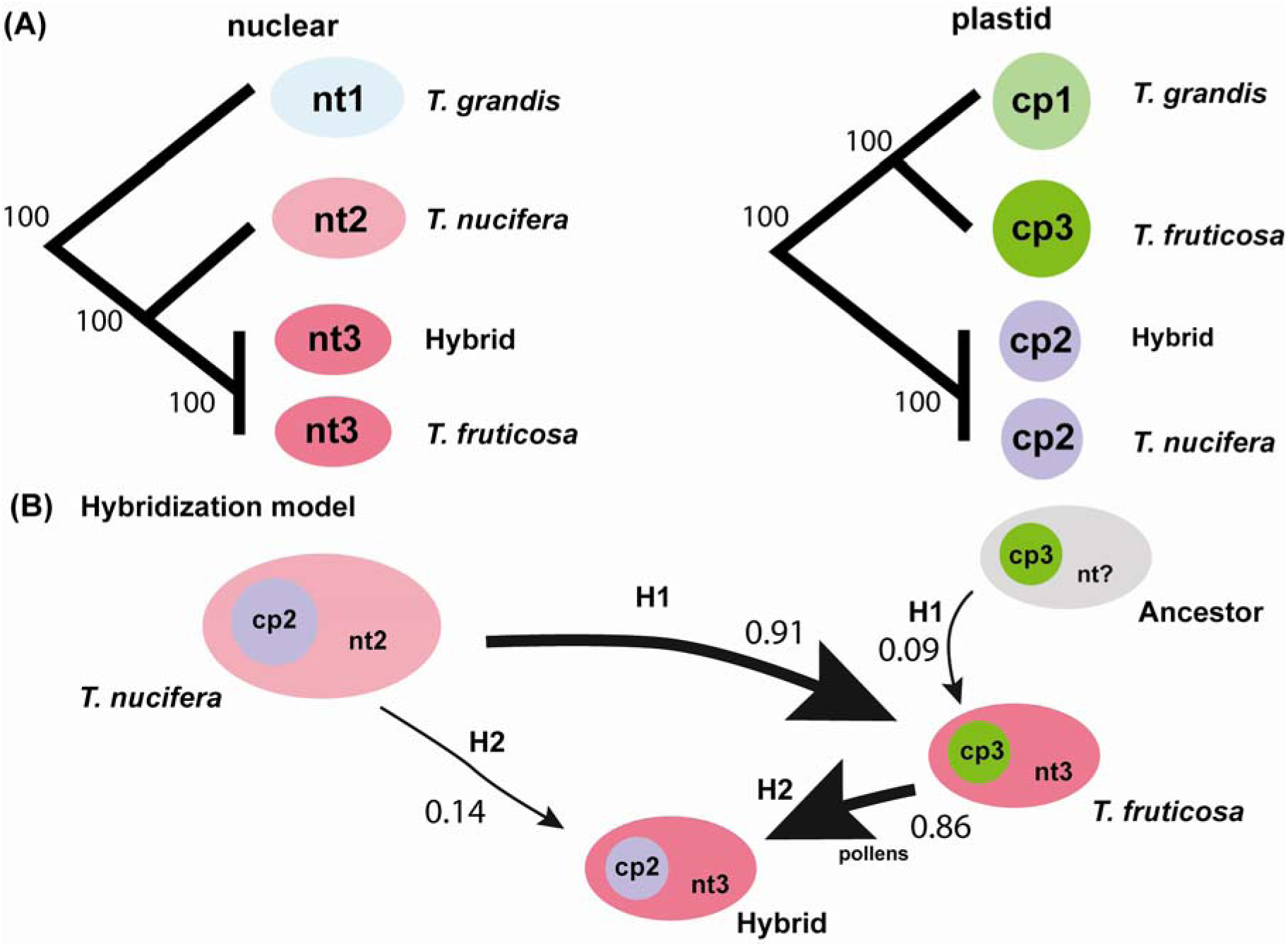
The hypothetical hybridization model of the formation of *Torreya fruticosa* might involve two plastid capture events. (A) cladogram of both nuclear and plastid tree among *T. grandis*, *T. nucifera*, *T. fruticosa*, and its Hybrid. (B) Hypothetical chloroplast capture events among *T. nucifera* and *T. fruticosa*, and their unknown ancestor suggest an ancient hybridization event (H1) between *T. nucifera* and an unidentified ancestral lineage, forming *T. fruticosa*. A more recent hybridization event (H2) between *T. nucifera* and *T. fruticosa* likely resulted in a mixed cytoplastome type in the putative hybrid. Numbers are indicating the percentage of genomes inherited from parents to descendants.

Due to the large overlapping distribution of *T. nucifera* and *T. fruticosa* (Fig. 2), the formation of hybrids may be prevented by pre-zygotic ecological barriers that have evolved as by-products of adaptation to their ecological niches (Baack et al., 2015; Coyne & Orr, 2004) (Fig. S1). For example, broadly sympatric species may still display ecological preferences for different habitats, e.g., available water source and soil microbes in *Boechera stricta* (Lee & Mitchell Olds, 2013; Wagner et al., 2015). Habitat modification, including increased disturbance and fragmentation, can erode ecological barriers and facilitate hybrid formation (Anderson, 1948; Buggs, 2007; Grabenstein & Taylor, 2018; Stebbins, 1950). This niche differentiation may further lead to different phenology of Japanese *Torreya*, including isolated flowering times as *Boechera* (Colautti et al., 2017). Alternatively, the limited formation of hybrids could result from the post-zygotic barriers, such as endosperm-based barrier (Abbott et al., 2013; Lafon Placette & Köhler, 2016). As this is the first time noticing the natural hybridization in Japanese *Torreya* (Taxaceae), we need to focus on the micro environmental and biological factors, such as soil, flowering time, and ploidy level in the future, to confirm and study the mechanism of this putative hybrid. Although we have detected many environmental factors related to precipitation and temperature have been differentiated between *T. fruticosa* and *T. nucifera* (Fig. S1), deeper studies on the hybrid zone are demanded.

### 4.2 Plastome evolution

The differentiated habitats (Fig. 2) of *T. nucifera* and *T. fruticosa* do not appear to significantly affect much on plastid evolution (Fig. 3). Since the full-length structure was not fully resolved by short reads, we are not able to compare the structures of plastomes in *Torreya*. Instead, we only focused on the gene mutations. Overall, we found the plastomes of *T. fruticosa* and its putative hybrid share the same gene number and gene order with other published *Torreya* species (Aizawa & Worth, 2021; Tao et al., 2016; Zhang et al., 2019). Different from the canonical quadripartite structure shared by most angiosperms plastomes (Daniell et al., 2016; Palmer & Stein, 1986), the plastomes of conifers, including our sequenced *Torreya* samples, are usually characterized with the loss of an IR (Fu et al., 2019; Hsu et al., 2014; Palmer & Thompson, 1982; Raubeson & Jansen, 1992; Wu et al., 2011; Wu & Chaw, 2014; Yi et al., 2013). In *T. fruticosa* and *T. nucifera*, we also found a trnQ-IR regions with 278 bp long (Fig. 3), which is congruent with discovery from Zhang *et al*. (2014) and Fu *et al*. (2019).

The only two substantially mutated plastid genes in *Torreya fruticosa* are *ycf1* and *accD*. Although the function of *ycf1* gene is still unknown, Dong et al. (2015) proposed that *ycf1* can be considered as important barcode due to their long length and high mutations, which might lead to important evolutionary functions. Regardless the absent *ycf1* gene in Poaceae (Goremykin et al., 2005), ycf1 has been applied to many phylogenetic works in Pinanceae (Gernandt et al., 2009), Orchidaceae (Neubig et al., 2009), Lamiaceae (Drew & Sytsma, 2012), and Annonaceae (Neubig & Abbott, 2010). The *accD* gene encodes acetyl-coenzyme A carboxylase, which is a multi-subunit protein complex that incorporates nuclear-encoded polypeptides and participates in fatty acid metabolism (Ohlrogge & Browse, 1995). The plastid *accD* encodes one subunit of the four-unit complex and has been reported as missing gene in numerous taxa (Ruhlman & Jansen, 2018), including e.g. *Trifolium* (Magee et al., 2010), *Juncus* (Zhou et al., 2023); Cyperaceae (*Cyperus*, Ren *et al*., 2021; *Elocharis*, (C. Lee et al., 2020). The 15 amino acid (aa) C-terminal catalytic domain of the *accD* protein, which is minimally required for prokaryotic ACCase function (S. S. Lee et al., 2004). Our study indicates that *ycf1* might involve at least six insertions/deletions in Japanese *Torreya* (Fig. S4) and *accD* gene in *T. fruticosa* had substantial 30-bp deletions when compared to *T. nucifera* and putative hybrid, leading to a decrease in the terminal length of *accD* gene that may impair its function (Fig. 3C). Different length of *ycf1* and *accD* gene have also been reported in Zhang *et al*. (2019). Our results support the expansion of *accD* gene (∼880 bp) in *Torreya* (Gymnosperms) compared to *Arabidopsis* (Angiosperms) (489 aa) (Sato et al., 1999). However, the validation of these two gene functions in *Torreya* needs further studies.

### 4.3 Application of whole genome sequencing

With the growing availability of genomic data from non-model systems (Chen et al., 2018; Marks et al., 2021), low-coverage whole genome sequencing (lc-WGS) has become an effective tool for phylogenomics and population genomics (Kumar et al., 2021; Martchenko & Shafer, 2023). However, for the species with large genome sizes, RAD-seq still might be one of the most convenient and cost-effective way to obtain the genetic information (Andrews et al., 2016; Barbanti et al., 2020; Wang et al., 2012). Although RAD-seq revolutionized population genomics by providing cost-effective SNP datasets without known reference data, it may cause later collected samples incompatible with previous RAD-seq data due to allelic dropout (ADO) by enzyme site mutations and libraries inconsistency. To address this ADO in continuous projects, our hybrid approach can combine the cost-efficiency of RAD-seq with the comprehensive genomic coverage of whole-genome sequencing (WGS) through RADADOR v.2, enabling phylogenetic reconstruction of samples from mixed datasets without any reference genomes. So far, lc-WGS data analysis remains more complex than RAD-seq tools (e.g. ipyrad and Stacks; Catchen et al., 2011), and lacks standardized pipelines to map to the well assembled reference (e.g. BWA, BWA-SW, SOAP2, bowtie, and bowtie2) and filter the low-quality loci (Picard, GATK) (Langmead et al., 2009; Langmead & Salzberg, 2012; Li & Durbin, 2009, 2010; Li et al., 2009; Van der Auwera & O’Connor, 2020). Our approach offers a more user-friendly alternative to traditional mapping methods. Specifically, GeneMiner, a k-mer based tool, allows for rapid and highly accurate identification of corresponding loci (Xie et al., 2024). Our study demonstrates the potential to integrate RAD-seq and WGS datasets for non-model species, which is particularly beneficial for individuals sampled at different times. In our case, given the large genome size of *Torreya* (∼20 Gb), we generated approximately 600 Gb of sequencing data for just six individuals. Nonetheless, this method would be even more efficient for non-model species with smaller or more typical genome sizes.

## Conclusions

We reclassified *Torreya nucifera* var. *radicans* Nakai as a species *T. fruticosa* Nakai and first-time revealed a potential natural hybridization zone in Japan *Torreya* based on genome-wide molecular data as well as their differentiated morphologies and ecological niches. All evidence supports a monophyletic group of *T. fruticosa* separated from *T. nucifera*. Our result suggested the formation of *T. fruticosa* might trace back to late Oligocene by an ancient hybridization event and the formation of the putative hybrid is more recent. This leads to a conflict between plastome phylogeny and nuclear phylogeny of Japanese *Torreya*. The putative hybrid individual has identical plastome with *T. nucifera* instead of *T. fruticosa*. Finally, we developed a pipeline (RADADOR v.2) first time addressing the allele dropout issue in continuous RAD-seq projects using low-coverage whole genome sequencing data.

## Acknowledgements

Dr. W. Zhou’s efforts on this project were supported in part by National Science Foundation Grant IOS-2034929 to Dr. Corbin Jones. This work is funded by Japan Society for the Promotion of Science KAKENHI Grant Number 24H00055, 24K02098, CAS-JSPS Bilateral Program of ‘Genomics of Biodiversity: Exploring Diversity Formation, Conservation and Utilization of Keystone Species in Sino-Japanese Floristic Region (SJFR)’. Special thanks to Drs. Soltis from University of Florida, Dr. Yan Yu from Sichuan University, and Dr. Paul Gabrielson, Dr. Jeremy Wang, Dr. Derick Poindexter, and Dr. Corbin Jones from University of North Carolina at Chapel Hill to read through the manuscript and provide valuable suggestions.

## Data Accessibility

All raw reads of six *Torreya fruiticosa* individuals were deposited in NCBI with Bioproject: PRJNA1276096, SRA accession numbers from SAMN49033941 to SAMN49033946, and GenBank IDs from PV874301 to PV874306. The rbcL of three *T. fruiticosa* individuals are available with GenBank IDs from PV867286 to PV867288; the *trnL*-*trnF* sequences are under GenBank IDs from PV867290 to PV867301; and all ITS sequences are with GenBank IDs from PV825776 to PV825787. The data matrices were deposited in Dryad (DOI: 10.5061/dryad.59zw3r2n1). And all code was deposited in Github (https://github.com/Bean061/RAD_Allele_Dropout_Remedy2).

## Author Contributions

Dr. Shota Sakaguchi recognized the taxa in the field and collected plant materials, conducted genotyping for all 16 samples and whole genome sequencing for two accessions. Besides, he read proof and polish the draft. Dr. Wenbin Zhou designed the experiment, developed the pipeline to retrieve shared RAD-seq loci from lcWGS data and conducted all data analyses, drafted the paper.

## Notes

### Competing Interest Statement

The authors have declared no competing interest.

## References

Abbott, R., Albach, D., Ansell, S., Arntzen, J. W., Baird, S. J. E., Bierne, N., Boughman, J., Brelsford, A., Buerkle, C. A., Buggs, R., Butlin, R. K., Dieckmann, U., Eroukhmanoff, F., Grill, A., Cahan, S. H., Hermansen, J. S., Hewitt, G., Hudson, A. G., Jiggins, C., … Zinner, D. (2013). Hybridization and speciation. Journal of Evolutionary Biology, 26(2), 229–246. 10.1111/j.1420-9101.2012.02599.x

Aizawa, M., & Worth, J. R. P. (2021). Phylogenetic origin of two Japanese Torreya taxa found in two regions with strongly contrasting snow depth. Journal of Plant Research, 134(5), 907–919. 10.1007/s10265-021-01301-8

Anderson, E. (1948). Hybridization of the habitat. Evolution, 1–9.

Andrews, K. R., Good, J. M., Miller, M. R., Luikart, G., & Hohenlohe, P. A. (2016). Harnessing the power of RADseq for ecological and evolutionary genomics. Nature Reviews Genetics, 17(2), 81–92.

Arnold, M. L. (1997). Natural hybridization and evolution. Oxford University Press.

Baack, E., Melo, M. C., Rieseberg, L. H., & Ortiz-Barrientos, D. (2015). The origins of reproductive isolation in plants. New Phytologist, 207(4), 968–984.

Barbanti, A., Torrado, H., Macpherson, E., Bargelloni, L., Franch, R., Carreras, C., & Pascual, M. (2020). Helping decision making for reliable and cost-effective 2b-RAD sequencing and genotyping analyses in non-model species. Molecular Ecology Resources, 20(3), 795–806.

Barrington, D. S., Haufler, C. H., & Werth, C. R. (1989). Hybridization, reticulation, and species concepts in the ferns. American Fern Journal, 79(2), 55–64.

Bernhardt, N., Brassac, J., Dong, X., Willing, E.-M., Poskar, C. H., Kilian, B., & Blattner, F. R. (2020). Genome-wide sequence information reveals recurrent hybridization among diploid wheat wild relatives. The Plant Journal, 102(3), 493–506.

Bouckaert, R., Heled, J., Kühnert, D., Vaughan, T., Wu, C.-H., Xie, D., Suchard, M. A., Rambaut, A., & Drummond, A. J. (2014). BEAST 2: A software platform for Bayesian evolutionary analysis. PLoS Computational Biology, 10(4), e1003537.

Buchholz, J. T. (1940). The embryogeny of Torreya, with a note on Austrotaxus. Bulletin of the Torrey Botanical Club, 731–754.

Buerkle, C. A., & Gompert, Z. (2013). Population genomics based on low coverage sequencing: How low should we go? Molecular Ecology, 22(11), 3028–3035.

Buggs, R. (2007). Empirical study of hybrid zone movement. Heredity, 99(3), 301–312.

Burke, J. (1975). Human use of the California nutmeg tree, Torreya californica, and of other members of the genus. Economic Botany, 127–139.

Capella-Gutiérrez, S., Silla-Martínez, J. M., & Gabaldón, T. (2009). trimAl: A tool for automated alignment trimming in large-scale phylogenetic analyses. Bioinformatics, 25(15), 1972–1973.

Catchen, J. M., Amores, A., Hohenlohe, P., Cresko, W., & Postlethwait, J. H. (2011). Stacks: Building and genotyping loci de novo from short-read sequences. G3: Genes| Genomes| Genetics, 1(3), 171–182.

Chen, F., Dong, W., Zhang, J., Guo, X., Chen, J., Wang, Z., Lin, Z., Tang, H., & Zhang, L. (2018). The sequenced angiosperm genomes and genome databases. Frontiers in Plant Science, 9, 418.

Chen, S., Zhou, Y., Chen, Y., & Gu, J. (2018). fastp: An ultra-fast all-in-one FASTQ preprocessor. Bioinformatics, 34(17), i884–i890.

Colautti, R. I., Ågren, J., & Anderson, J. T. (2017). Phenological shifts of native and invasive species under climate change: Insights from the *Boechera–Lythrum* model. Philosophical Transactions of the Royal Society B: Biological Sciences, 372(1712), 20160032. 10.1098/rstb.2016.0032

Conkle, M. T., & Critchfield, W. B. (1988). Genetic variation and hybridization of ponderosa pine. In: Ponderosa Pine: The Species and Its Management, Washington State University Cooperative Extension, 1988: P. 27-43.

Coyne, J. A., & Orr, H. A. (2004). Speciation: A catalogue and critique of species concepts. Philosophy of Biology: An Anthology, 272–292.

Currat, M., Ruedi, M., Petit, R. J., & Excoffier, L. (2008). The hidden side of invasions: Massive introgression by local genes. Evolution, 62(8), 1908–1920.

Daniell, H., Lin, C.-S., Yu, M., & Chang, W.-J. (2016). Chloroplast genomes: Diversity, evolution, and applications in genetic engineering. Genome Biology, 17, 1–29.

Darling, A. C., Mau, B., Blattner, F. R., & Perna, N. T. (2004). Mauve: Multiple alignment of conserved genomic sequence with rearrangements. Genome Research, 14(7), 1394–1403.

Darzentas, N. (2010). Circoletto: Visualizing sequence similarity with Circos. Bioinformatics, 26(20).

Davey, J. W., Cezard, T., Fuentes-Utrilla, P., Eland, C., Gharbi, K., & Blaxter, M. L. (2013). Special features of RAD Sequencing data: Implications for genotyping. Molecular Ecology, 22(11), 3151–3164.

De La Harpe, M., Paris, M., Karger, D. N., Rolland, J., Kessler, M., Salamin, N., & Lexer, C. (2017). Molecular ecology studies of species radiations: Current research gaps, opportunities and challenges. Molecular Ecology, 26(10), 2608–2622. 10.1111/mec.14110

Dong, W., Xu, C., Li, C., Sun, J., Zuo, Y., Shi, S., Cheng, T., Guo, J., & Zhou, S. (2015). Ycf1, the most promising plastid DNA barcode of land plants. Scientific Reports, 5(1), 8348. 10.1038/srep08348

Donoghue, M. J., & Smith, S. A. (2004). Patterns in the assembly of temperate forests around the Northern Hemisphere. Philosophical Transactions of the Royal Society of London. Series B: Biological Sciences, 359(1450), 1633–1644.

Dowling, T. E., & DeMarais, B. D. (1993). Evolutionary significance of introgressive hybridization in cyprinid fishes. Nature, 362(6419), 444–446.

Doyle, J. (1991). DNA Protocols for Plants. In G. M. Hewitt, A. W. B. Johnston, & J. P. W. Young (Eds.), Molecular Techniques in Taxonomy (pp. 283–293). Springer Berlin Heidelberg. 10.1007/978-3-642-83962-7_18

Drew, B. T., & Sytsma, K. J. (2012). Phylogenetics, biogeography, and staminal evolution in the tribe Mentheae (Lamiaceae). American Journal of Botany, 99(5), 933–953.

Eaton, D. A., & Overcast, I. (2020). ipyrad: Interactive assembly and analysis of RADseq datasets. Bioinformatics, 36(8), 2592–2594.

Eckenwalder, J. E. (1984). Natural intersectional hybridization between North American species of Populus (Salicaceae) in sections Aigeiros and Tacamahaca. II. Taxonomy. Canadian Journal of Botany, 62(2), 325–335.

Eckenwalder, J. E. (2009). Conifers of the world: The complete reference. Timber press.

eFloras (2008). Published on the Internet http://www.efloras.org [accessed 7 August 2025] Missouri Botanical Garden, St. Louis, MO & Harvard University Herbaria, Cambridge, MA.

Fishman, L., & Willis, J. H. (2001). Evidence for Dobzhansky-Muller incompatibilites contributing to the sterility of hybrids between Mimulus guttatus and M. nasutus. Evolution, 55(10), 1932–1942.

Frankel, L. E., & Ané, C. (2023). Summary tests of introgression are highly sensitive to rate variation across lineages. Systematic Biology, 72(6), 1357–1369.

Fu, C.-N., Wu, C.-S., Ye, L.-J., Mo, Z.-Q., Liu, J., Chang, Y.-W., Li, D.-Z., Chaw, S.-M., & Gao, L.-M. (2019). Prevalence of isomeric plastomes and effectiveness of plastome super-barcodes in yews (Taxus) worldwide. Scientific Reports, 9(1), 2773.

Gautier, M., Gharbi, K., Cezard, T., Foucaud, J., Kerdelhué, C., Pudlo, P., Cornuet, J.-M., & Estoup, A. (2013). The effect of RAD allele dropout on the estimation of genetic variation within and between populations. Molecular Ecology, 22(11), 3165–3178.

Genovart, M. (2009). Natural hybridization and conservation. Biodiversity and Conservation, 18(6), 1435–1439. 10.1007/s10531-008-9550-x

Gernandt, D. S., Hernández-León, S., Salgado-Hernández, E., & Pérez de La Rosa, J. A. (2009). Phylogenetic relationships of Pinus subsection Ponderosae inferred from rapidly evolving cpDNA regions. Systematic Botany, 34(3), 481–491.

Goremykin, V. V., Holland, B., Hirsch-Ernst, K. I., & Hellwig, F. H. (2005). Analysis of Acorus calamus chloroplast genome and its phylogenetic implications. Molecular Biology and Evolution, 22(9), 1813–1822.

Grabenstein, K. C., & Taylor, S. A. (2018). Breaking Barriers: Causes, Consequences, and Experimental Utility of Human-Mediated Hybridization. Trends in Ecology & Evolution, 33(3), 198–212. 10.1016/j.tree.2017.12.008

Grant, P. R., & Grant, B. R. (1992). Hybridization of bird species. Science, 256(5054), 193–197.

Grant, V., & Grant, K. A. (1971). Natural hybridization between the cholla cactus species Opuntia spinosior and Opuntia versicolor. Proceedings of the National Academy of Sciences, 68(9), 1993–1995.

Hijmans, R. J., Van Etten, J., Cheng, J., Mattiuzzi, M., Sumner, M., Greenberg, J. A., Lamigueiro, O. P., Bevan, A., Racine, E. B., Shortridge, A., & others. (2015). Package ‘raster.’ R Package, 734, 473.

Hils, MH. (1996). Taxaceae. In: Flora of North America Editorial Committee eds. Flora of North America north of Mexico [Online]. 21. New York and Oxford. Vol. 2. Available from http://www.efloras.org/florataxon.aspx?flora_id=1%26taxon_id=10871 [accessed 08 June 2021]

Hohenlohe, P. A., Funk, W. C., & Rajora, O. P. (2021). Population genomics for wildlife conservation and management. Molecular Ecology, 30(1), 62–82. 10.1111/mec.15720

Hsu, C.-Y., Wu, C.-S., & Chaw, S.-M. (2014). Ancient nuclear plastid DNA in the yew family (Taxaceae). Genome Biology and Evolution, 6(8), 2111–2121.

Huson, D. H., & Bryant, D. (2006). Application of phylogenetic networks in evolutionary studies. Molecular Biology and Evolution, 23(2), 254–267.

Jin, J.-J., Yu, W.-B., Yang, J.-B., Song, Y., DePamphilis, C. W., Yi, T.-S., & Li, D.-Z. (2020). GetOrganelle: A fast and versatile toolkit for accurate de novo assembly of organelle genomes. Genome Biology, 21, 1–31.

Johnson, M. G., Pokorny, L., Dodsworth, S., Botigué, L. R., Cowan, R. S., Devault, A., Eiserhardt, W. L., Epitawalage, N., Forest, F., Kim, J. T., & others. (2019). A universal probe set for targeted sequencing of 353 nuclear genes from any flowering plant designed using k-medoids clustering. Systematic Biology, 68(4), 594–606.

Katoh, K., & Standley, D. M. (2013). MAFFT multiple sequence alignment software version 7: Improvements in performance and usability. Molecular Biology and Evolution, 30(4), 772–780.

Kearse, M., Moir, R., Wilson, A., Stones-Havas, S., Cheung, M., Sturrock, S., Buxton, S., Cooper, A., Markowitz, S., Duran, C., & others. (2012). Geneious Basic: An integrated and extendable desktop software platform for the organization and analysis of sequence data. Bioinformatics, 28(12), 1647–1649.

Kou, Y.-X., Xiao, K., Lai, X.-R., Wang, Y.-J., & Zhang, Z.-Y. (2017). Natural hybridization between Torreya jackii and T. grandis (Taxaceae) in southeast China. Journal of Systematics and Evolution, 55(1), 25–33.

Krutovskii, K. V., & Bergmann, F. (1995). Introgressive hybridization and phylogenetic relationships between Norway, Picea abies (L.) Karst., and Siberian, P. obovata Ledeb., spruce species studied by isozyme loci. Heredity, 74(5), 464–480.

Kumar, P., Choudhary, M., Jat, B., Kumar, B., Singh, V., Kumar, V., Singla, D., & Rakshit, S. (2021). Skim sequencing: An advanced NGS technology for crop improvement. Journal of Genetics, 100, 1–10.

Lafon Placette, C., & Köhler, C. (2016). Endosperm based postzygotic hybridization barriers: Developmental mechanisms and evolutionary drivers. Molecular Ecology, 25(11), 2620–2629. 10.1111/mec.13552

Lagache, L., Klein, E. K., Guichoux, E., & Petit, R. J. (2013). Fine-scale environmental control of hybridization in oaks. Molecular Ecology, 22(2), 423–436.

Langmead, B., & Salzberg, S. L. (2012). Fast gapped-read alignment with Bowtie 2. Nature Methods, 9(4), 357–359.

Langmead, B., Trapnell, C., Pop, M., & Salzberg, S. L. (2009). Ultrafast and memory-efficient alignment of short DNA sequences to the human genome. Genome Biology, 10(3), R25.

Leaché, A. D., & Oaks, J. R. (2017). The utility of single nucleotide polymorphism (SNP) data in phylogenetics. Annual Review of Ecology, Evolution, and Systematics, 48(1), 69–84.

Lee, C., & Mitchell Olds, T. (2013). Complex trait divergence contributes to environmental niche differentiation in ecological speciation of *B oechera stricta*. Molecular Ecology, 22(8), 2204–2217. 10.1111/mec.12250

Lee, C., Ruhlman, T. A., & Jansen, R. K. (2020). Unprecedented intraindividual structural heteroplasmy in Eleocharis (Cyperaceae, Poales) plastomes. Genome Biology and Evolution, 12(5), 641–655.

Lee, S. S., Jeong, W. J., Bae, J. M., Bang, J. W., Liu, J. R., & Harn, C. H. (2004). Characterization of the plastid-encoded carboxyltransferase subunit (accD) gene of potato. Molecules and Cells, 17(3), 422–429.

Li, H., & Durbin, R. (2009). Fast and accurate short read alignment with Burrows–Wheeler transform. Bioinformatics, 25(14), 1754–1760.

Li, H., & Durbin, R. (2010). Fast and accurate long-read alignment with Burrows–Wheeler transform. Bioinformatics, 26(5), 589–595.

Li, J., Milne, R. I., Ru, D., Miao, J., Tao, W., Zhang, L., Xu, J., Liu, J., & Mao, K. (2020). Allopatric divergence and hybridization within Cupressus chengiana (Cupressaceae), a threatened conifer in the northern Hengduan Mountains of western China. Molecular Ecology, 29(7), 1250–1266.

Li, R., Yu, C., Li, Y., Lam, T.-W., Yiu, S.-M., Kristiansen, K., & Wang, J. (2009). SOAP2: An improved ultrafast tool for short read alignment. Bioinformatics, 25(15), 1966–1967.

Lipman, M. J., Chester, M., Soltis, P. S., & Soltis, D. E. (2013). Natural hybrids between Tragopogon mirus and T. miscellus (Asteraceae): A new perspective on karyotypic changes following hybridization at the polyploid level. American Journal of Botany, 100(10), 2016–2022.

Lou, H., Song, L., Li, X., Zi, H., Chen, W., Gao, Y., Zheng, S., Fei, Z., Sun, X., & Wu, J. (2023). The Torreya grandis genome illuminates the origin and evolution of gymnosperm-specific sciadonic acid biosynthesis. Nature Communications, 14(1), 1315.

Magee, A. M., Aspinall, S., Rice, D. W., Cusack, B. P., Sémon, M., Perry, A. S., Stefanović, S., Milbourne, D., Barth, S., Palmer, J. D., & others. (2010). Localized hypermutation and associated gene losses in legume chloroplast genomes. Genome Research, 20(12), 1700–1710.

Malinsky, M., Trucchi, E., Lawson, D.J. and Falush, D. (2018). RADpainter and fineRADstructure: population inference from RADseq data. Molecular biology and evolution, 35(5), pp.1284–1290.

Mamanova, L., Coffey, A. J., Scott, C. E., Kozarewa, I., Turner, E. H., Kumar, A., Howard, E., Shendure, J., & Turner, D. J. (2010). Target-enrichment strategies for next-generation sequencing. Nature Methods, 7(2), 111–118.

Marks, R. A., Hotaling, S., Frandsen, P. B., & VanBuren, R. (2021). Representation and participation across 20 years of plant genome sequencing. Nature Plants, 7(12), 1571–1578.

Martchenko, D., & Shafer, A. B. (2023). Contrasting whole-genome and reduced representation sequencing for population demographic and adaptive inference: An alpine mammal case study. Heredity, 131(4), 273–281.

Mastretta-Yanes, A., Arrigo, N., Alvarez, N., Jorgensen, T. H., Piñero, D., & Emerson, B. C. (2015). Restriction site-associated DNA sequencing, genotyping error estimation and de novo assembly optimization for population genetic inference. Molecular Ecology Resources, 15(1), 28–41.

Meleshko, O., Martin, M. D., Korneliussen, T. S., Schröck, C., Lamkowski, P., Schmutz, J., Healey, A., Piatkowski, B. T., Shaw, A. J., Weston, D. J., & others. (2021). Extensive genome-wide phylogenetic discordance is due to incomplete lineage sorting and not ongoing introgression in a rapidly radiated bryophyte genus. Molecular Biology and Evolution, 38(7), 2750–2766.

Menon, M., Bagley, J. C., Page, G. F., Whipple, A. V., Schoettle, A. W., Still, C. J., Wehenkel, C., Waring, K. M., Flores-Renteria, L., Cushman, S. A., & others. (2021). Adaptive evolution in a conifer hybrid zone is driven by a mosaic of recently introgressed and background genetic variants. Communications Biology, 4(1), 160.

Minh, B. Q., Schmidt, H. A., Chernomor, O., Schrempf, D., Woodhams, M. D., Von Haeseler, A., & Lanfear, R. (2020). IQ-TREE 2: New models and efficient methods for phylogenetic inference in the genomic era. Molecular Biology and Evolution, 37(5), 1530–1534.

Mo, Z.-Q., Wang, J., Möller, M., Yang, J.-B., & Gao, L.-M. (2023). Phylogenetic relationships and next-generation barcodes in the Genus Torreya reveal a high proportion of misidentified cultivated plants. International Journal of Molecular Sciences, 24(17), 13216.

Neubig, K. M., & Abbott, J. R. (2010). Primer development for the plastid region ycf1 in Annonaceae and other magnoliids. American Journal of Botany, 97(6), e52–e55.

Neubig, K. M., Whitten, W. M., Carlsward, B. S., Blanco, M. A., Endara, L., Williams, N. H., & Moore, M. (2009). Phylogenetic utility of ycf 1 in orchids: A plastid gene more variable than mat K. Plant Systematics and Evolution, 277(1), 75–84.

North, H. L., McGaughran, A., & Jiggins, C. D. (2021). Insights into invasive species from whole-genome resequencing. Molecular Ecology, 30(23), 6289–6308.

Ohashi, H. (2015). Taxaceae in Wild Flowers of Japan, Revised Edition vol.1, p42–44 (pp. 392), Edited by Ojashi, H., Kadota, Y., Murata, J., Yonekura, K.

Ohlrogge, J., & Browse, J. (1995). Lipid biosynthesis. The Plant Cell, 7(7), 957.

Ōi, Jisaburō. (1965). Flora of Japan. Smithsonian Institution. Retrieved from https://library.si.edu/digital-library/book/floraofjapaninen00oiji

Ou, Q., Huang, X., Pan, D., Wang, S., Huang, Y., Lu, S., Wang, Y., & Kou, Y. (2025). The Divergence History of Two Japanese Torreya Taxa (Taxaceae): Implications for Species Diversification in the Japanese Archipelago. Plants, 14(10), 1537. 10.3390/plants14101537

Palmer, J. D., & Stein, D. B. (1986). Conservation of chloroplast genome structure among vascular plants. Current Genetics, 10, 823–833.

Palmer, J. D., & Thompson, W. F. (1982). Chloroplast DNA rearrangements are more frequent when a large inverted repeat sequence is lost. Cell, 29(2), 537–550.

Payseur, B. A., & Rieseberg, L. H. (2016). A genomic perspective on hybridization and speciation. Molecular Ecology, 25(11), 2337–2360.

Petit, R. J., Bodénès, C., Ducousso, A., Roussel, G., & Kremer, A. (2004). Hybridization as a mechanism of invasion in oaks. New Phytologist, 161(1), 151–164.

Picard toolkit. (2019). In Broad Institute, GitHub repository. Broad Institute. https://broadinstitute.github.io/picard/

Raubeson, L. A., & Jansen, R. K. (1992). A rare chloroplast-DNA structural mutation is shared by all conifers. Biochemical Systematics and Ecology, 20(1), 17–24.

Ren, W., Guo, D., Xing, G., Yang, C., Zhang, Y., Yang, J., Niu, L., Zhong, X., Zhao, Q., Cui, Y., & others. (2021). Complete chloroplast genome sequence and comparative and phylogenetic analyses of the cultivated Cyperus esculentus. Diversity, 13(9), 405.

Rieseberg, L. H., & Carney, S. E. (1998). Plant hybridization. The New Phytologist, 140(4), 599–624.

Rieseberg, L. H., Raymond, O., Rosenthal, D. M., Lai, Z., Livingstone, K., Nakazato, T., Durphy, J. L., Schwarzbach, A. E., Donovan, L. A., & Lexer, C. (2003). Major ecological transitions in wild sunflowers facilitated by hybridization. Science, 301(5637), 1211–1216.

Ruhlman, T. A., & Jansen, R. K. (2018). Aberration or analogy? The atypical plastomes of Geraniaceae. In Advances in botanical research (Vol. 85, pp. 223–262). Elsevier.

Sato, S., Nakamura, Y., Kaneko, T., Asamizu, E., & Tabata, S. (1999). Complete structure of the chloroplast genome of Arabidopsis thaliana. DNA Research, 6(5), 283–290.

Seehausen, O. (2004). Hybridization and adaptive radiation. Trends in Ecology & Evolution, 19(4), 198–207.

Shin, S., Kim, S.-C., Hong, K. N., Kang, H., & Lee, J.-W. (2019). The complete chloroplast genome of Torreya nucifera (Taxaceae) and phylogenetic analysis. Mitochondrial DNA Part B, 4(2), 2537–2538.

Soltis, P. S., & Soltis, D. E. (2009). The Role of Hybridization in Plant Speciation. Annual Review of Plant Biology, 60(1), 561–588. 10.1146/annurev.arplant.043008.092039

Song, B., Ning, W., Wei, D., Jiang, M., Zhu, K., Wang, X., Edwards, D., Odeny, D. A., & Cheng, S. (2023). Plant genome resequencing and population genomics: Current status and future prospects. Molecular Plant, 16(8), 1252–1268. 10.1016/j.molp.2023.07.009

Stalter, R., & Dial, S. (1984). Environmental status of the stinking cedar, Torreya taxifolia. Bartonia, 50, 40–42.

Stebbins, G. L. (1950). Variation and evolution in plants. Columbia University Press.

Stull, G. W., Qu, X.-J., Parins-Fukuchi, C., Yang, Y.-Y., Yang, J.-B., Yang, Z.-Y., Hu, Y., Ma, H., Soltis, P. S., Soltis, D. E., Li, D.-Z., Smith, S. A., & Yi, T.-S. (2021). Gene duplications and phylogenomic conflict underlie major pulses of phenotypic evolution in gymnosperms. Nature Plants, 7(8), 1015–1025. 10.1038/s41477-021-00964-4

Tao, K., Gao, L., Li, J., Chen, S., Su, Y., & Wang, T. (2016). The complete chloroplast genome of Torreya fargesii (Taxaceae). Mitochondrial DNA Part A, 27(5), 3512–3513.

Taylor, S. A., & Larson, E. L. (2019). Insights from genomes into the evolutionary importance and prevalence of hybridization in nature. Nature Ecology & Evolution, 3(2), 170–177.

Than, C., Ruths, D., & Nakhleh, L. (2008). PhyloNet: A software package for analyzing and reconstructing reticulate evolutionary relationships. BMC Bioinformatics, 9, 1–16.

Tillich, M., Lehwark, P., Pellizzer, T., Ulbricht-Jones, E. S., Fischer, A., Bock, R., & Greiner, S. (2017). GeSeq–versatile and accurate annotation of organelle genomes. Nucleic Acids Research, 45(W1), W6–W11.

Tsuda, Y., Chen, J., Stocks, M., Källman, T., Sønstebø, J. H., Parducci, L., Semerikov, V., Sperisen, C., Politov, D., Ronkainen, T., & others. (2016). The extent and meaning of hybridization and introgression between Siberian spruce (Picea obovata) and Norway spruce (Picea abies): Cryptic refugia as stepping stones to the west? Molecular Ecology, 25(12), 2773–2789.

Twyford, A. D., & Ennos, R. A. (2012). Next-generation hybridization and introgression. Heredity, 108(3), 179–189. 10.1038/hdy.2011.68

Untergasser, A., Cutcutache, I., Koressaar, T., Ye, J., Faircloth, B. C., Remm, M., & Rozen, S. G. (2012). Primer3—New capabilities and interfaces. Nucleic Acids Research, 40(15), e115–e115.

Vallejo Marín, M., & Hiscock, S. J. (2016). Hybridization and hybrid speciation under global change. New Phytologist, 211(4), 1170–1187. 10.1111/nph.14004

Van der Auwera, G. A., & O’Connor, B. D. (2020). Genomics in the cloud: Using Docker, GATK, and WDL in Terra. O’Reilly Media.

Wagner, M. R., Lundberg, D. S., Coleman Derr, D., Tringe, S. G., Dangl, J. L., & Mitchell□Olds, T. (2015). Corrigendum to Wagner *et al*.: Natural soil microbes alter flowering phenology and the intensity of selection on flowering time in a wild Arabidopsis relative. Ecology Letters, 18(2), 218–220. 10.1111/ele.12400

Wang, S., Meyer, E., McKay, J. K., & Matz, M. V. (2012). 2b-RAD: a simple and flexible method for genome-wide genotyping. Nature Methods, 9(8), 808–810.

Wen, J., Ickert-Bond, S., Nie, Z.-L., & Li, R. (2010). Timing and modes of evolution of eastern Asian-North American biogeographic disjunctions in seed plants. Darwin’s Heritage Today: Proceedings of the Darwin 2010 Beijing International Conference, 252–269.

Wen, J., Nie, Z.-L., & Ickert-Bond, S. M. (2016). Intercontinental disjunctions between eastern Asia and western North America in vascular plants highlight the biogeographic importance of the Bering land bridge from late Cretaceous to Neogene. Journal of Systematics and Evolution, 54(5), 469–490.

Wick, R. R., Schultz, M. B., Zobel, J., & Holt, K. E. (2015). Bandage: Interactive visualization of de novo genome assemblies. Bioinformatics, 31(20), 3350–3352.

Wickham, H. (2011). Ggplot2. Wiley Interdisciplinary Reviews: Computational Statistics, 3(2), 180–185.

Worth, J. R., Larcombe, M. J., Sakaguchi, S., Marthick, J. R., Bowman, D. M., Ito, M., & Jordan, G. J. (2016). Transient hybridization, not homoploid hybrid speciation, between ancient and deeply divergent conifers. American Journal of Botany, 103(2), 246–259.

Wu, C.-S., & Chaw, S.-M. (2014). Highly rearranged and size-variable chloroplast genomes in conifers II clade (cupressophytes): Evolution towards shorter intergenic spacers. Plant Biotechnology Journal, 12(3), 344–353.

Wu, C.-S., Lin, C.-P., Hsu, C.-Y., Wang, R.-J., & Chaw, S.-M. (2011). Comparative chloroplast genomes of Pinaceae: Insights into the mechanism of diversified genomic organizations. Genome Biology and Evolution, 3, 309–319.

Wu, S., Wang, Y., Wang, Z., Shrestha, N., & Liu, J. (2022). Species divergence with gene flow and hybrid speciation on the Qinghai–Tibet Plateau. New Phytologist, 234(2), 392–404.

Xie, P., Guo, Y., Teng, Y., Zhou, W., & Yu, Y. (2024). GeneMiner: A tool for extracting phylogenetic markers from next-generation sequencing data. Molecular Ecology Resources, 24(3), e13924.

Yamazaki, T. (1995) Taxaceae. In: Iwatsuki K, Yamazaki, T., Boufford, D.E., Ohba, H. (eds) Flora of Japan, vol 1. Kodansha, Tokyo, pp 286–287.

Yi, X., Gao, L., Wang, B., Su, Y.-J., & Wang, T. (2013). The complete chloroplast genome sequence of Cephalotaxus oliveri (Cephalotaxaceae): Evolutionary comparison of Cephalotaxus chloroplast DNAs and insights into the loss of inverted repeat copies in gymnosperms. Genome Biology and Evolution, 5(4), 688–698.

Zhang, X., Zhang, H.-J., Landis, J. B., Deng, T., Meng, A.-P., Sun, H., Peng, Y.-S., Wang, H.-C., & Sun, Y.-X. (2019). Plastome phylogenomic analysis of Torreya (Taxaceae). Journal of Systematics and Evolution, 57(6), 607–615.

Zhang, Y., Ma, J., Yang, B., Li, R., Zhu, W., Sun, L., Tian, J., & Zhang, L. (2014). The complete chloroplast genome sequence of Taxus chinensis var. Mairei (Taxaceae): Loss of an inverted repeat region and comparative analysis with related species. Gene, 540(2), 201–209.

Zhou, W., Armijos, C. E., Lee, C., Lu, R., Wang, J., Ruhlman, T. A., Jansen, R. K., Jones, A. M., & Jones, C. D. (2023). Plastid Genome Assembly Using Long-read data. Molecular Ecology Resources, 23(6), 1442–1457. 10.1111/1755-0998.13787

Zhou, W., Harris, A., & Xiang, Q. (Jenny). (2022). Phylogenomics and biogeography of *Torreya* (Taxaceae)—Integrating data from three organelle genomes, morphology, and fossils and a practical method for reducing missing data from RAD seq. Journal of Systematics and Evolution, 60(6), 1241–1262. 10.1111/jse.12838

Zhou, W., & Xiang, Q.-Y. J. (2022). Phylogenomics AND biogeography of Castanea (chestnut) and Hamamelis (witch-hazel)–Choosing between RAD-seq and Hyb-Seq approaches. Molecular Phylogenetics and Evolution, 176, 107592.

Zhou, Y. F., Abbott, R. J., Jiang, Z. Y., Du, F. K., Milne, R. I., & Liu, J. Q. (2010). Gene flow and species delimitation: A case study of two pine species with overlapping distributions in southeast China. Evolution, 64(8), 2342–2352.

