## Supplementary figures for "Bypassing the Batch Effects to Improve the Reusability of RAD-seq Data: A Case Study of Recurrent Hybridization in the Japanese *Torreya* Species Complex"

### Supplementary figure legend

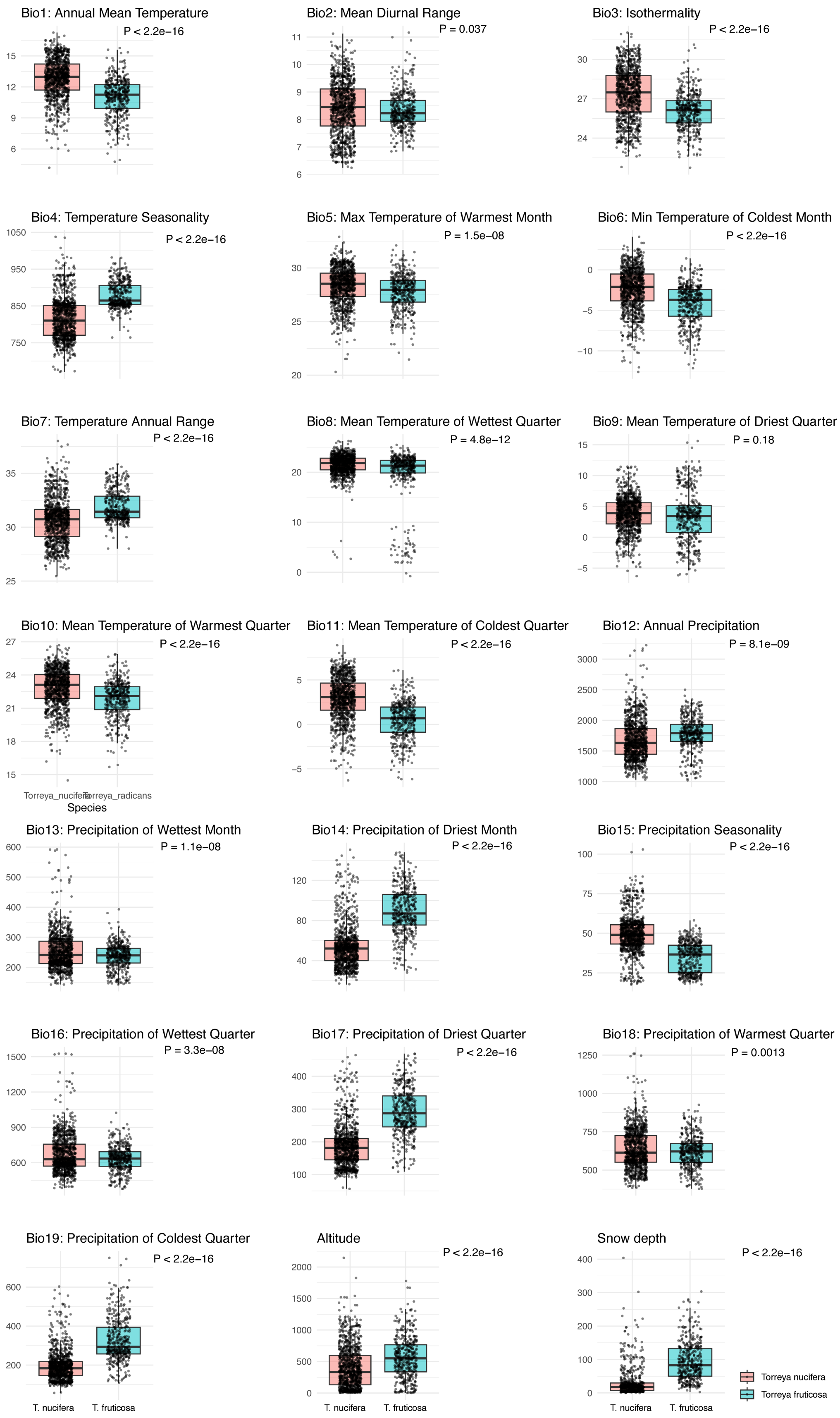

Bio1: Annual Mean Temperature
P < 2.2e−16
Bio2: Mean Diurnal Range
P = 0.037
Bio3: Isothermality
P < 2.2e−16
Bio4: Temperature Seasonality
P < 2.2e−16
Bio5: Max Temperature of Warmest Month
P = 1.5e−08
Bio6: Min Temperature of Coldest Month
P < 2.2e−16
Bio7: Temperature Annual Range
P < 2.2e−16
Bio8: Mean Temperature of Wettest Quarter
P = 4.8e−12
Bio9: Mean Temperature of Driest Quarter
P = 0.18
Bio10: Mean Temperature of Warmest Quarter
P < 2.2e−16
Torreya_nucifera
Torreya_radicans
Species
Bio11: Mean Temperature of Coldest Quarter
P < 2.2e−16
Bio12: Annual Precipitation
P = 8.1e−09
Bio13: Precipitation of Wettest Month
P = 1.1e−08
Bio14: Precipitation of Driest Month
P < 2.2e−16
Bio15: Precipitation Seasonality
P < 2.2e−16
Bio16: Precipitation of Wettest Quarter
P = 3.3e−08
Bio17: Precipitation of Driest Quarter
P < 2.2e−16
Bio18: Precipitation of Warmest Quarter
P = 0.0013
Bio19: Precipitation of Coldest Quarter
P < 2.2e−16
T. nucifera
T. fruticosa
Altitude
P < 2.2e−16
Snow depth
P < 2.2e−16
Torreya nucifera
Torreya fruticosa


Figure S1.

Climatic niche comparison between *Torreya nucifera* and *T. fruticosa*. A total of 20 bioclimatic variables were obtained from WorldClim, including 11 temperature-related and 8 precipitation-related layers, and one snow depth layer were obtained from https://nlftp.mlit.go.jp/ksj/gml/datalist/KsjTmplt-G02-v3_0.html. A total of 19 out of 21 climate layers indicates significant difference between Japanese *Torreya* species complex. Only Bio2 (Mean Diurnal Range) and Bio9 (Mean Temparature of Driest Quarter) are not differentiated.

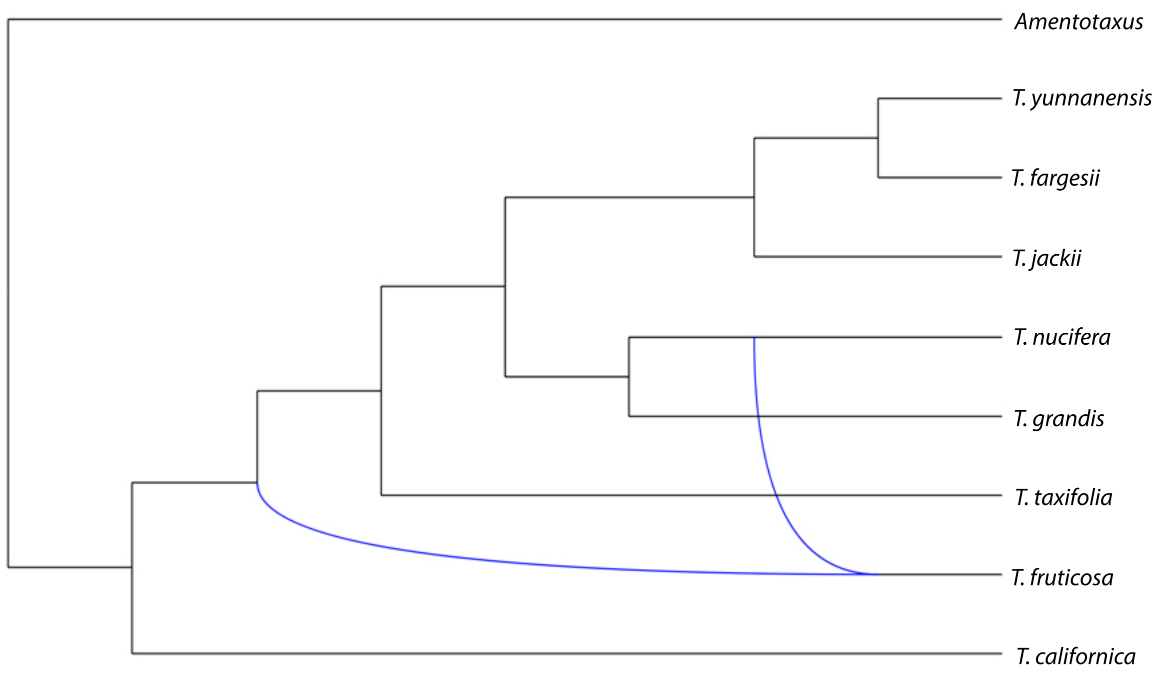


Figure S2. PhyloNet analysis of *Torreya* given the assumption that only one hybridization event occurred among the evolutionary history of *Torreya*.

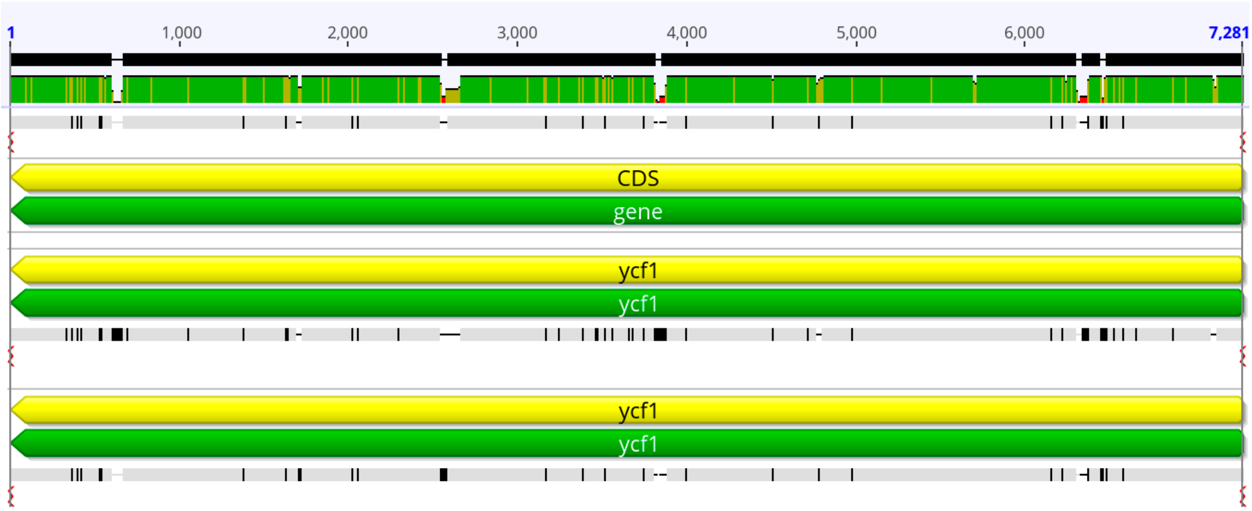

Figure S3. Mutation of ycf1 gene among Japanese *Torreya* species complex, including *T. nucifera* in the 1st line, *T. fruticosa* in the 2nd line, and putative Hybrid B in the 3rd line.
